# Polycomb repressive complex 1.1 coordinates homeostatic and emergency myelopoiesis

**DOI:** 10.1101/2022.09.07.507016

**Authors:** Yaeko Nakajima-Takagi, Motohiko Oshima, Junichiro Takano, Shuhei Koide, Naoki Itokawa, Shun Uemura, Masayuki Yamashita, Shohei Andoh, Kazumasa Aoyama, Yusuke Isshiki, Daisuke Shinoda, Atsunori Saraya, Fumio Arai, Kiyoshi Yamaguchi, Yoichi Furukawa, Haruhiko Koseki, Tomokatsu Ikawa, Atsushi Iwama

## Abstract

Polycomb repressive complex (PRC) 1 regulates stem cell fate by mediating mono-ubiquitination of histone H2A at lysine 119. While canonical PRC1 is critical for hematopoietic stem and progenitor cell (HSPC) maintenance, the role of non-canonical PRC1 in hematopoiesis remains elusive. PRC1.1, a non-canonical PRC1, consists of PCGF1, RING1B, KDM2B, and BCOR. We recently showed that PRC1.1 insufficiency induced by the loss of PCGF1 or BCOR causes myeloid-biased hematopoiesis and promotes transformation of hematopoietic cells in mice. Here we show that PRC1.1 serves as an epigenetic switch that coordinates homeostatic and emergency hematopoiesis. PRC1.1 maintains balanced output of steady-state hematopoiesis by restricting C/EBPα-dependent precocious myeloid differentiation of HSPCs and the HOXA9- and β-catenin-driven self-renewing network in myeloid progenitors. Upon regeneration, PRC1.1 is transiently inhibited to facilitate formation of granulocyte-macrophage progenitor (GMP) clusters, thereby promoting emergency myelopoiesis. Moreover, constitutive inactivation of PRC1.1 results in unchecked expansion of GMPs and eventual transformation. Collectively, our results define PRC1.1 as a novel critical regulator of emergency myelopoiesis, dysregulation of which leads to myeloid transformation.

## INTRODUCTION

While lifelong hematopoiesis is considered driven by hematopoietic stem cells (HSCs)(Sawai et al., 2016; Säwen et al., 2018), recent evidence pointed out a major role for multipotent progenitors (MPPs) and lineage-committed progenitors in hematopoiesis (Busch et al., 2015; Sun et al., 2014). Hematopoietic stem and progenitor cells (HSPCs) are highly responsive to various stresses such as infection, inflammation, and myeloablation (Trumpp et al., 2010; Zhao & Baltimore, 2015), which facilitate myelopoiesis by activating HSPCs to undergo precocious myeloid differentiation and transiently amplifying myeloid progenitors that rapidly differentiate into mature myeloid cells (Hérault et al., 2017; Pietras et al., 2016). This reprogramming of HSPCs, termed “emergency myelopoiesis”, serves to immediately replenish mature myeloid cells to control infection and regeneration (Manz & Boettcher, 2014). Recent evidence further suggested that uncontrolled activation of the myeloid regeneration programs results in the development of chronic inflammatory diseases and hematological malignancies (Chiba et al., 2018; Zhao & Baltimore, 2015). Emergency myelopoiesis is driven via activation of key myeloid transcriptional networks at the HSPC and myeloid progenitor cell levels (Rosenbauer & Tenen, 2007). However, the epigenetic regulatory mechanisms governing emergency myelopoiesis remained largely unknown.

Polycomb group (PcG) proteins are the key epigenetic regulators of a variety of biological processes (Piunti & Shilatifard, 2021). They comprise the multiprotein complexes, polycomb repressive complex (PRC) 1 and PRC2, which establish and maintain the transcriptional repression through histone modifications. PRC1 and PRC2 add mono-ubiquitination at lysine 119 of histone H2A (H2AK119ub) and mono-, di-, and tri-methylation at lysine 27 of histone H3 (H3K27me1/me2/me3), respectively, and cooperatively repress transcription (Blackledge et al., 2015; Iwama, 2017). PRC1 complexes are divided into subgroups (PRC1.1 to PRC1.6) according to the subtype of the Polycomb group ring finger (PCGF) subunits (PCGF1-6). PCGF2/MEL18 and PCGF4/BMI1 act as components of canonical PRC1 (PRC1.2 and 1.4, respectively) that are recruited to its target sites in a manner dependent on H3K27me3, whereas the others (PCGF1, 3, 5, and 6) constitute non-canonical PRC1 (PRC1.1, PRC1.3, PRC1.5, and PRC1.6, respectively) that are recruited independently of H3K27me3 (Blackledge et al., 2014; Gao et al., 2012; Wang et al., 2004).

PCGF4/BMI1-containing canonical PRC1 (PRC1.4) has been characterized for its role in maintaining self-renewal capacity and multipotency of HSCs (Sashida & Iwama, 2012). We and others have reported that BMI1 transcriptionally represses the loci for *CDKN2A* and developmental regulator genes (e.g., B cell regulators) to maintain self-renewal capacity and multipotency of HSPCs (Iwama et al., 2004; Oguro et al., 2010; Park et al., 2003). We also reported that PCGF5-containing PRC1.5 regulates global levels of H2AK119ub, but is dispensable for HSPC function (Si et al., 2016). On the other hand, we and others recently showed that PRC1.1 components, PCGF1, KDM2B, and BCOR, maintain normal hematopoiesis and suppress malignant transformation of hematopoietic cells (Andricovich et al., 2016; Isshiki et al., 2019; Tara et al., 2018). PRC1.1 consists of PCGF1, RING1A/B, KDM2B, and BCOR or BCLRL1. KDM2B binds to non-methylated CpG islands through its DNA-binding domain, thereby recruiting other PRC1.1 components (Farcas et al., 2012; He et al., 2013). PCGF1 was found to restrict the proliferative capacity of myeloid progenitor cells by down-regulating *Hoxa* family genes in *in vitro* knockdown experiments (Ross et al., 2012). Correspondingly, we demonstrated that Pcgf1 loss induces myeloid-biased hematopoiesis and promotes JAK2V617F-induced myelofibrosis in mice (Shinoda *et al*, 2022). Bcor loss also showed myeloid-biased hematopoiesis and promoted the initiation and progression of myelodysplastic syndrome in collaboration with Tet2 loss (Tara *et al*, 2018). However, detailed analysis of the role for PRC1.1 in hematopoiesis, especially in the context of hematopoietic regeneration and emergency myelopoiesis, have not yet been reported.

Here, we analyzed the murine hematopoiesis in the absence of PCGF1 and uncovered critical roles of PCGF1-containing PRC1.1 in homeostatic, emergency, and malignant hematopoiesis.

## RESULTS

### PCGF1 restricts myeloid commitment of HSPCs

To understand the function of PCGF1 in hematopoiesis, we crossed *Pcgf1^fl^* mice, in which exons 2 to 7 are floxed (Fig S1A), with *Rosa26::Cre-ERT2* (*Cre-ERT*) mice (*Cre-ERT;Pcgf1^fl/fl^*). To delete *Pcgf1* specifically in hematopoietic cells, we transplanted bone marrow (BM) cells from *Cre-ERT* control and *Cre-ERT;Pcgf1^fl/fl^* CD45.2 mice into lethally irradiated CD45.1 recipient mice and deleted *Pcgf1* by intraperitoneal injection of tamoxifen (Figs 1A and S1B). We confirmed the efficient deletion of *Pcgf1* in donor-derived hematopoietic cells from the peripheral blood (PB) by genomic PCR (Fig S1C). RT-qPCR confirmed the significant reduction of *Pcgf1* mRNA lacking exons 2 to 7 in donor-derived BM Lineage marker^−^Sca-1^+^c-Kit^+^ (LSK) HSPCs (Fig S1D). We hereafter refer to the recipient mice reconstituted with *Cre-ERT* control and *Cre-ERT;Pcgf1^fl/fl^* cells treated with tamoxifen as control and *Pcgf1^Δ/Δ^* mice, respectively.

**Figure 1.**
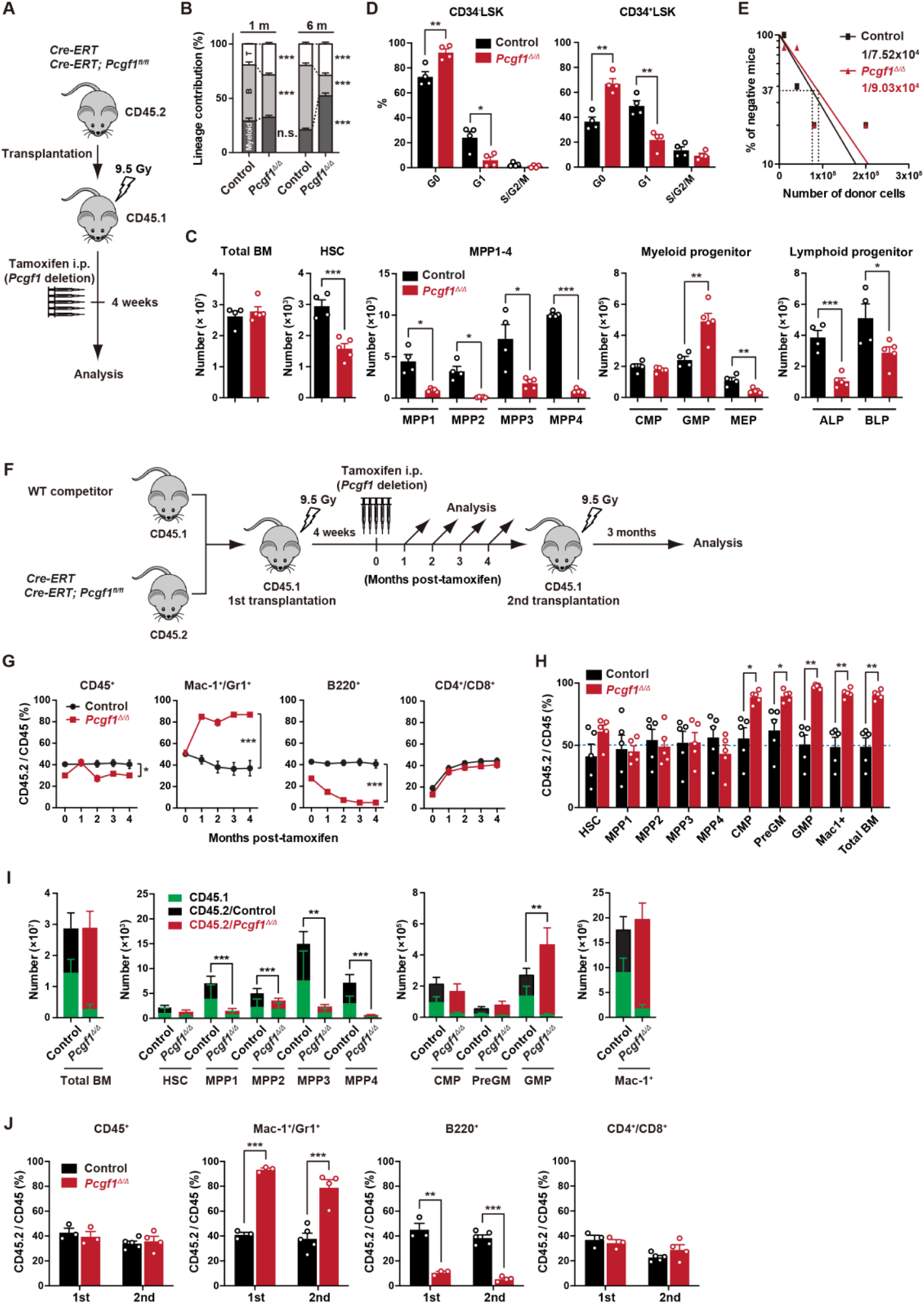
PCGF1 regulates myelopoiesis but not self-renewal of HSPCs. (A) Strategy for analyzing *Pcgf1^Δ/Δ^* hematopoietic cells. Total BM cells (5×10^6^) from *Cre-ERT* and *Cre-ERT;Pcgf1^fl/fl^* were transplanted into lethally irradiated CD45.1 recipient mice. *Pcgf1* was deleted by intraperitoneal injections of tamoxifen at 4 weeks post-transplantation. (B) The proportions of Mac-1^+^ and/or Gr-1^+^ myeloid cells, B220^+^ B cells, and CD4^+^ or CD8^+^ T cells among CD45.2^+^ donor-derived hematopoietic cells in the PB from control (n=9) and *Pcgf1^Δ/Δ^*(n=14) mice. (C) Absolute numbers of total BM cells, HSCs, MPPs, myeloid progenitors, and CLPs (ALP and BLP) in a unilateral pair of femur and tibia 4 weeks after the tamoxifen injection (n=4-5). (D) Cell cycle status of CD34^−^LSK HSCs and CD34^+^LSK MPPs assessed by Ki67 and 7-AAD staining 4 weeks after the tamoxifen injection. (E) *In vivo* limiting dilution assay. Limiting numbers of BM cells (1×10^4^, 4×10^4^, 8×10^4^, and 2×10^5^) isolated from BM of primary recipients (control and *Pcgf1^Δ/Δ^* mice after transplantation) were transplanted into sublethally irradiated secondary recipient mice with 2×10^5^ of competitor CD45.1 BM cells (n=5 each). Due to the low contribution of *Pcgf1^Δ/Δ^* HSPCs to B cells, mice with chimerism of donor myeloid and T cells more than 1% in the PB at 16 weeks after transplantation were considered to be engrafted successfully, and the others were defined as non-engrafted mice. The frequencies of HSPCs that contributed to both myeloid and T cells are indicated. (F) Strategy for analyzing *Pcgf1^Δ/Δ^* hematopoietic cells. Total BM cells (2×10^6^) from *Cre-ERT* and *Cre-ERT;Pcgf1^fl/fl^*CD45.2 mice were transplanted into lethally irradiated CD45.1 recipient mice with the same number of competitor CD45.1 BM cells. *Pcgf1* was deleted by intraperitoneal injections of tamoxifen at 4 weeks post-transplantation. Secondary transplantation was performed using 5×10^6^ total BM cells from primary recipients at 4 months post intraperitoneal injections of tamoxifen. (G) The chimerism of CD45.2 donor cells in PB CD45^+^ leukocytes, Mac-1^+^ and/or Gr1^+^ myeloid cells, B220^+^ B cells, and CD4^+^ or CD8^+^ T cells in control and *Pcgf1^Δ/Δ^* mice (n=6 each) after the tamoxifen injection. (H) The chimerism of CD45.2 donor-derived cells in BM 4 weeks after the tamoxifen injection (n=5). (I) Absolute numbers of CD45.1 and CD45.2 total BM cells, HSCs, MPPs, myeloid progenitors, and Mac-1^+^ mature myeloid cells in a unilateral pair of femur and tibia 4 weeks after the tamoxifen injection (n=5). (J) The chimerism of CD45.2 donor-derived cells in PB in primary (n=3 each) and secondary (n=4-5) transplantation. Data are shown as the mean ± SEM. \**p*<0.05, \*\**p*<0.01, \*\*\**p*<0.001 by the Student’s *t*-test. Each symbol is derived from an individual mouse. A representative of more than two independent experiments is shown.

*Pcgf1^Δ/Δ^* mice exhibited mild anemia and leukopenia, which was mainly attributed to the reduction in B-cell numbers (Fig 1B). Myeloid cell numbers in PB were relatively maintained and their proportion increased over time in *Pcgf1^Δ/Δ^* mice (Figs 1B and S2A). While BM cellularity was comparable between control and *Pcgf1^Δ/Δ^* mice, mature myeloid cells were increased at the expense of B cells in *Pcgf1^Δ/Δ^* BM (Figs 1C, S2B and C). Among the committed progenitor cells, the numbers of granulocyte-macrophage progenitors (GMPs) were significantly increased in *Pcgf1^Δ/Δ^* BM, whereas those of megakaryocyte-erythroid progenitors (MEPs), pre- and pro-B cells, all-lymphoid progenitors (ALPs) and B-cell-biased lymphoid progenitors (BLPs) were decreased (Figs 1C, S2B and 3B). Of interest, *Pcgf1^Δ/Δ^* mice showed reduction in the numbers of HSCs and all subsets of MPPs in BM (Figs 1C and S3A). The reduction in the HSPC pool size was accompanied by decreased cells in cycling phases in *Pcgf1^Δ/Δ^* HSPCs (CD34^−^ and CD34^+^ LSK cells) (Fig 1D). Despite the reduction in phenotypic HSPCs, limiting dilution assays with competitive BM transplantation revealed that the numbers of HSPCs that established long-term repopulation of myeloid and T cells (B cells were excluded due to the low contribution of *Pcgf1^Δ/Δ^* HSPCs to B cells, see below) were comparable between control and *Pcgf1^Δ/Δ^*BM (Fig. 1E). Extramedullary hematopoiesis was evident in the *Pcgf1^Δ/Δ^* spleen, as the absolute numbers of HSPCs, GMPs, and mature myeloid cells were markedly increased (Figs S2B, D and S3C). Differentiation of thymocytes in the *Pcgf1^Δ/Δ^* thymus was largely normal (Fig S2B).

To further evaluate the role of PCGF1 in hematopoiesis, we transplanted BM cells with the same number of CD45.1 wild-type (WT) competitor cells (Fig 1F). In this competitive setting, only a mild decrease was detected in the overall chimerism of CD45.2^+^ *Pcgf1^Δ/Δ^*cells in PB (Fig 1G). In contrast, the chimerism of *Pcgf1^Δ/Δ^* cells in myeloid cells (Mac-1^+^ and/or Gr1^+^) was markedly increased while that in B cell lineage (B220^+^) was decreased. (Fig 1G). In BM, *Pcgf1^Δ/Δ^*cells outcompeted the competitor cells in the myeloid lineage compartments from the common myeloid progenitor (CMP) stage (Fig 1H). Since *Pcgf1^Δ/Δ^* cell showed reductions in the numbers of HSCs and MPPs in a non-competitive setting (Fig 1C), we examined the absolute numbers of test and competitor cells in this competitive repopulation. Of interest, the competitive *Pcgf1^Δ/Δ^* recipients also exhibit similar changes in BM hematopoietic cell numbers. Both CD45.2^+^ *Pcgf1^Δ/Δ^* and CD45.1^+^ WT cells were depleted in HSPC fractions in *Pcgf1^Δ/Δ^* recipients, while the total numbers of myeloid progenitors and mature myeloid cells were maintained or rather increased (Fig. 1I). These findings suggest that *Pcgf1^Δ/Δ^* hematopoietic cells suppress hematopoiesis driven by co-existing WT HSPCs through non-autonomous mechanisms. To evaluate the impact of PCGF1 loss on long-term hematopoiesis, we harvested BM cells from primary recipient mice 4 months after tamoxifen injection and transplanted them into secondary recipients. *Pcgf1^Δ/Δ^*cells reproduced the myeloid-biased hematopoiesis in secondary recipients (Fig. 1J).

We next evaluated the capacity of *Pcgf1^Δ/Δ^* HSCs to differentiate to myeloid and lymphoid cells in culture. *Pcgf1^Δ/Δ^*HSCs displayed slower growth under HSPC-expanding culture conditions (Fig S4A), which is in good agreement with fewer cycling *Pcgf1^Δ/Δ^* HSPCs (Fig 1D). Nevertheless, *Pcgf1^Δ/Δ^* HSCs showed better growth than control cells under myeloid culture conditions (Fig S4A). On the other hand, limiting dilution assays using a co-culture system with TSt-4 stromal cells (Masuda et al., 2005) revealed that the capacity of *Pcgf1^Δ/Δ^* HSCs to produce B and T cells was declined by 2- and 5-fold, respectively, compared to the control (Fig S4B). The discrepancy in T cell production between *in vitro* and *in vivo* may be due to the compensatory expansion of T cells in the thymus. These results indicate that PCGF1 loss enhances myelopoiesis at the expense of lymphopoiesis. These phenotypes were similar to those of mice expressing a carboxyl-terminal truncated BCOR that cannot interact with PCGF1 (Tara et al., 2018).

### PCGF1 inhibits precocious myeloid commitment of HSPCs through repression of C/EBPα, which is critical for balanced output of HSPCs

To clarify the molecular mechanisms underlying myeloid-biased differentiation of *Pcgf1^Δ/Δ^* HSPCs, we performed RNA sequence (RNA-seq) analysis of HSPCs from mice 4 weeks after the tamoxifen injection. Principal component analysis (PCA) showed shifts of the transcriptomic profiles of *Pcgf1^Δ/Δ^*MPP2 and MPP3 toward pre-GMs (Fig 2A). Gene set enrichment analysis (GSEA) revealed up-regulation of genes involved in myeloid cell development (Brown et al., 2006), CEBP targets (Gery et al., 2005), and genes up-regulated upon C/EBPα overexpression (Loke et al., 2018) in *Pcgf1^Δ/Δ^* HSPCs compared to controls (Fig 2B and Table S1). C/EBP family are master transcription factors for myeloid differentiation (Avellino & Delwel, 2017; Rosenbauer & Tenen, 2007). RT-PCR analysis demonstrated a significant up-regulation of *Cepba* in HSPCs, but not in GMPs (Fig 2C). C/EBPα drives myelopoiesis and antagonizes lymphoid differentiation (Fukuchi et al., 2006; Rosenbauer & Tenen, 2007; Xie et al., 2004). Disruption of *Cebpa* blocks the transition from common myeloid progenitors (CMPs) to GMPs (Zhang et al., 1997). So, we evaluated the contribution of de-repressed *Cebpa* in *Pcgf1^Δ/Δ^* HSPCs. First, we overexpressed *Cebpa* in WT LSK cells and cultured them on TSt-4 stromal cells. Only two-fold upregulation of *Cebpa* was sufficient to enhance myeloid differentiation and suppress B cell differentiation of HSPCs (Fig 2D). Conversely, the reduction of the levels of *Cebpa* expression by introducing a *Cebpa^fl/+^* allele in *Pcgf1^Δ/Δ^* LSK cells was sufficient to restore balanced production of myeloid and B cells by *Pcgf1^Δ/Δ^* HSPCs (Fig 2E). These results demonstrate that de-repression of *Cebpa* largely accounts for the myeloid-biased differentiation of *Pcgf1^Δ/Δ^* HSPCs.

**Figure 2.**
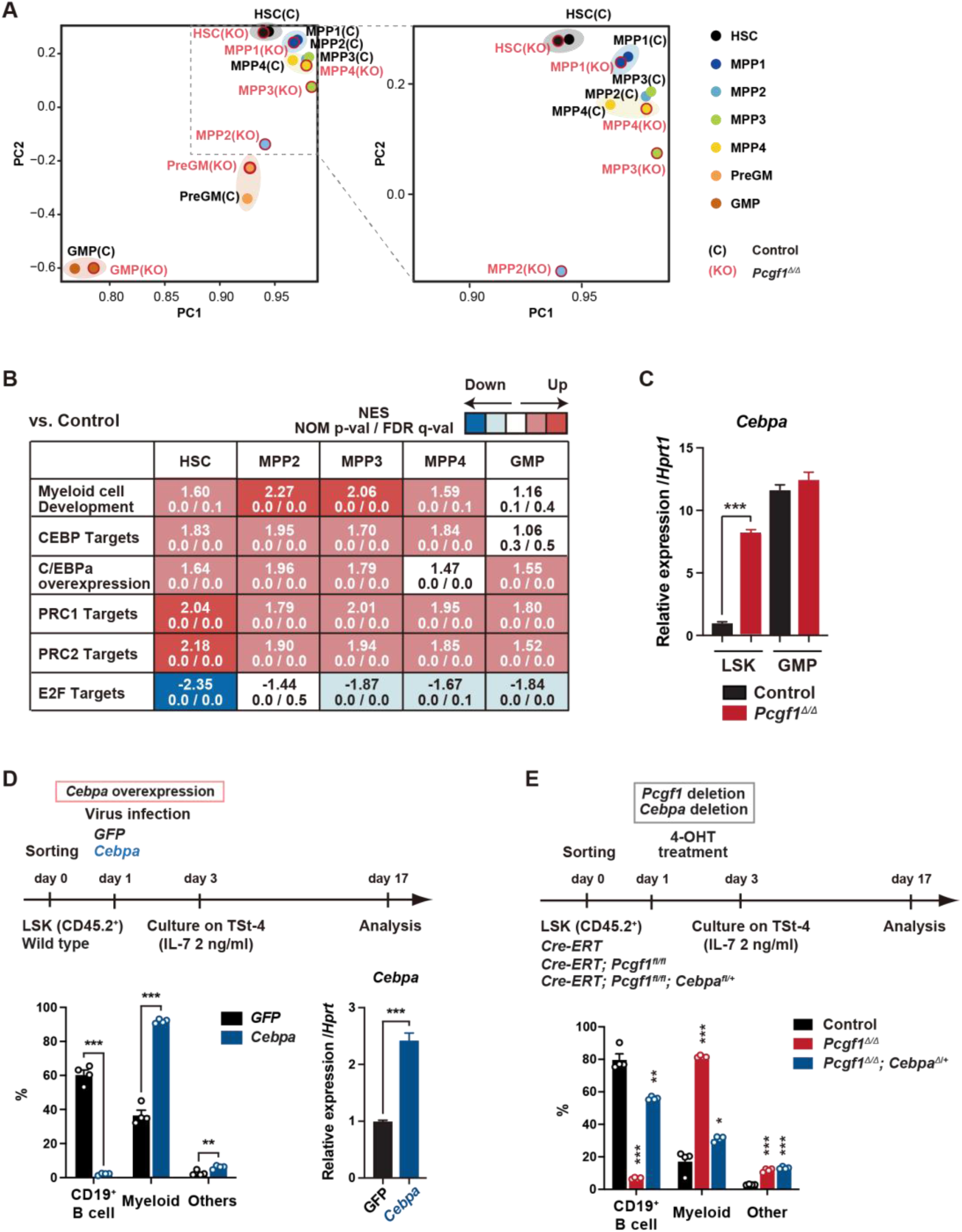
*Pcgf1*-deficient HSPCs undergo myeloid reprogramming. (A) Principal component (PC) analyses based on total gene expression obtained by RNA-seq of HSCs, MPPs, pre-GM, and GMPs from control and *Pcgf1^Δ/Δ^* mice. Magnified view of the boxed part is depicted on the right. (B) GSEA using RNA-seq data. Summary of GSEA data of representative gene sets is shown. Normalized enrichment scores (NES), nominal p values (NOM), and false discovery rates (FDR) are indicated. The gene sets used are indicated in Supplementary Table 1. (C) Quantitative RT-PCR analysis of *Cebpa* in LSK cells and GMPs. *Hprt1* was used to normalize the amount of input RNA. Data are shown as the mean ± SEM (n=3). (D) Effects of *Cebpa* overexpression on HSPC differentiation *in vitro*. LSK cells were transduced with either Control (*GFP*) or *Cebpa* retrovirus, then cultured on TSt-4 stromal cells in the presence of IL-7 (upper). The proportions of myeloid (Mac1^+^ and /or Gr-1^+^), B cells (CD19^+^), and others (Mac1^-^Gr-1^-^CD19^-^) among CD45.2^+^GFP^+^ hematopoietic cells on day17 of culture are indicated (lower left; n=4 each). RT-qPCR analysis of *Cebpa* in LSK cells transduced with control or *Cebpa* retrovirus on day 14 of culture (n=3). *Hprt1* was used to normalize the amount of input RNA (lower right). Each symbol is derived from an individual culture. (E) Impact of *Cebpa* haploinsufficiency on myeloid-biased differentiation of *Pcgf1^Δ/Δ^* HSPCs. LSK cells from *Cre-ERT*, *Cre-ERT;Pcgf1^fl/fl^*and *Cre-ERT;Pcgf1^fl/fl^;Cebpa^fl/+^* mice were treated with 4-OHT (200 nM) for 2 days in culture to delete *Pcgf1* and *Cebpa*. Cells were further cultured on TSt-4 stromal cells in the presence of IL-7 (upper). The proportions of myeloid (Mac1^+^ and /or Gr-1^+^), B cells (CD19^+^), and others (Mac1^-^Gr-1^-^CD19^-^) among CD45.2^+^GFP^+^ hematopoietic cells on day17 of culture are indicated (lower; n=4 each ; *, versus control). Each symbol is derived from an individual culture. \**p*<0.05, \*\**p*<0.01, \*\*\**p*<0.001 by the Student’s *t*-test.

We next attempted to rescue the myeloid-biased differentiation of *Pcgf1^Δ/Δ^* HSPCs by exogenous *Pcgf1* or a canonical PRC1 gene *Bmi1/Pcgf4*. We transduced *Cre-ERT;Pcgf1^fl/fl^* HSPCs to induce *Pcgf1* or *Bmi1* expression, transplanted them into lethally irradiated mice, and deleted endogenous *Pcgf1* (Fig S5B). The myeloid skew in *Pcgf1^Δ/Δ^* PB leukocytes was completely prevented by ectopic expression of *Pcgf1* but not of *Bmi1* (Fig S5B), highlighting distinct roles of non-canonical PRC1.1 and canonical PRC1 in hematopoietic differentiation.

We also noticed that the E2F targets (Ishida et al., 2001) were downregulated in *Pcgf1^Δ/Δ^* HSPCs (Fig 2B), which may underlie the disturbed cell cycle progression and delayed proliferation observed in *Pcgf1^Δ/Δ^*HSPCs (Figs S3D and S4A). C/EBPα represses E2F-mediated transcription (D’Alo’ et al., 2003; Slomiany et al., 2000), inhibits HSC cell cycle entry (Ye et al., 2013; Zhang et al., 2004), and promotes precocious IL-1β-driven emergency myelopoiesis (Higa et al., 2021). Thus, up-regulated *Cebpa* upon PCGF1 loss may inhibit cell cycle and promote myeloid commitment of HSPCs.

### Deletion of *Pcgf1* affects levels of H2AK119ub1

To understand how PCGF1 loss affects H2AK119ub1 status in HSPCs, we performed chromatin immunoprecipitation followed by sequencing (ChIP-seq) analysis using control and *Pcgf1^Δ/Δ^* HSPCs. Since none of the anti-PCGF1 antibodies were suitable for ChIP analysis, we used 3×Flag-PCGF1-expressing BM LK cells obtained by retrovirally transducing LSK cells and transplanting them to lethally irradiated mice. We defined “PRC1 targets” and “PRC2 targets” as genes with H2AK119ub1 and H3K27me3 enrichment greater than twofold over the input signals in control LSK cells at promoter regions [transcription start site (TSS) ± 2.0 kb], respectively (Table S2). GSEA revealed that both PRC1 and PRC2 targets were upregulated in *Pcgf1^Δ/Δ^* HSPCs and GMPs (Fig 2B). K-means clustering divided PRC1 targets into 2 clusters depending on the levels of H2AK119ub1 and H3K27me3. Cluster 1 genes (1835 RefSeq ID genes) were marked with high levels of H2AK119ub1 and H3K27me3 at promoter regions, while cluster 2 genes (2691 RefSeq ID genes) showed moderate levels of H2AK119ub1 and H3K27me3 (Fig 3A). PCGF1 showed stronger binding to the promoters of cluster 2 genes than cluster 1 genes (Fig 3A). Interestingly, only cluster 2 genes showed moderate but significant reductions in H2AK119ub1 and H3K27me3 levels in *Pcgf1^Δ/Δ^* HSPCs (Fig 3B). The loss of PCGF1 was also significantly associated with de-repression of PRC1 target genes in clusters 1 and 2 (Fig 3B). These results suggest that cluster 2 genes are the major targets of PCGF1-containing PRC1.1 and *Cebpa* was included in cluster 2 genes. Bivalent genes, which are enriched for developmental regulator genes marked with both active and repressive histone marks (H3K4me3 and H3K27me3, respectively, mostly with H2AK119ub1), are classical targets of canonical PRC1 and PRC2 (Bernstein et al., 2006; Ku et al., 2008) and are implicated in multipotency of HSPCs (Oguro et al., 2010). Of note, bivalent genes defined by our previous ChIP-seq data of HSPCs (Aoyama et al., 2018) (Table S2) were more enriched in cluster 1 genes than cluster 2 genes (Fig 3C).

**Figure 3.**
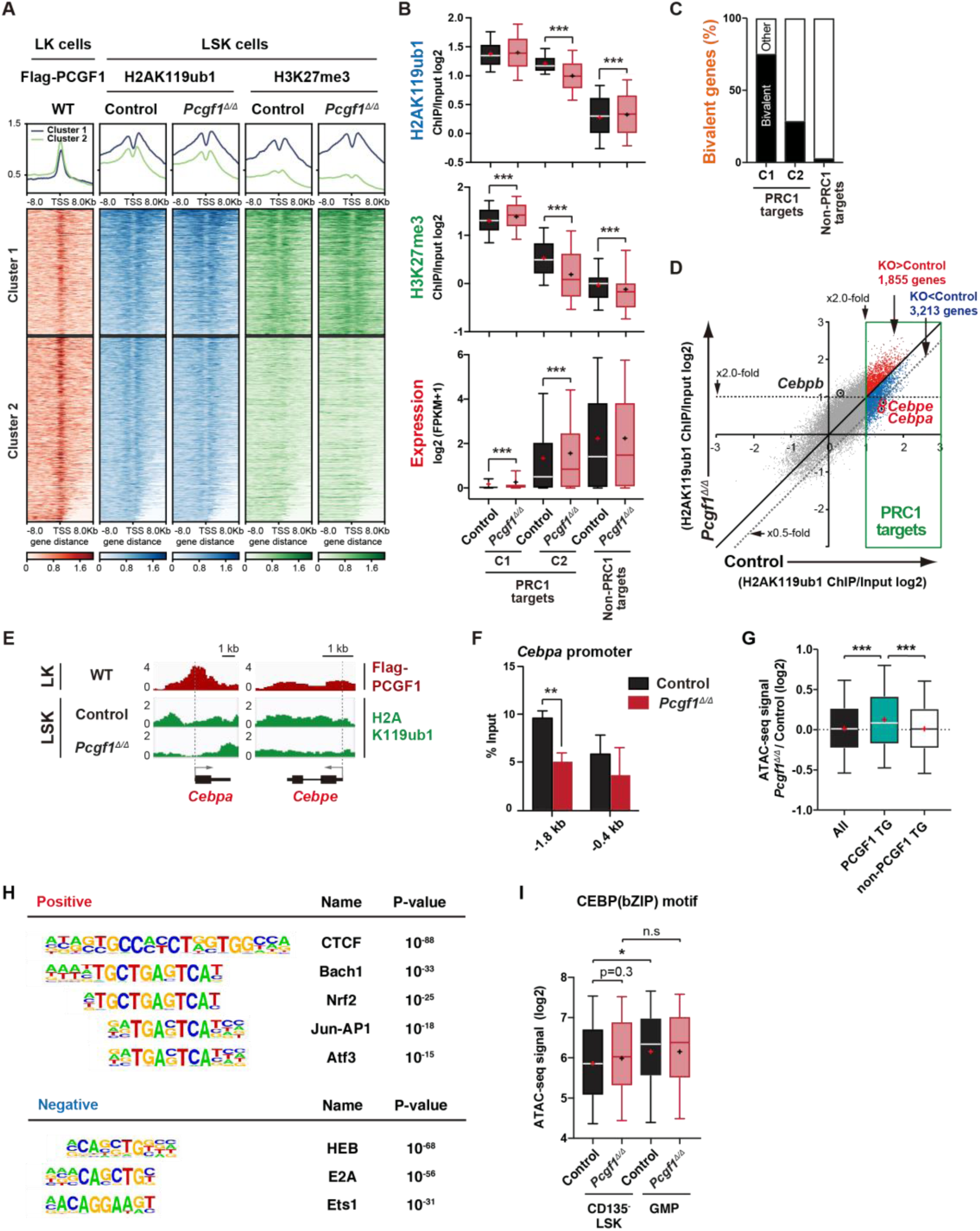
PCGF1 regulates local H2AK119ub1 levels in HSPCs. (A) K-means clustering of 3×FLAG-PCGF1, H2AK119ub1, and H3K27me3 ChIP peaks around TSS (± 8.0 kb) of PRC1 target genes. The average levels of ChIP peaks in each cluster are plotted in upper columns. (B) Box-and-whisker plots showing H2AK119ub1, H3K27me3, and transcription levels of genes in PRC1 targets (clusters 1 and 2) and non-PRC1 targets in control and *Pcgf1^Δ/Δ^* LSK cells. Boxes represent 25–75 percentile ranges. The whiskers represent 10–90 percentile ranges. Horizontal bars represent medians. Mean values are indicated by “+”. (C) Proportion of bivalent genes in PRC1 targets (clusters 1 and 2) and non-PRC1 targets in LSK cells. Bivalent genes were defined using our previous ChIP-seq data of wild-type LSK cells (Aoyama et al., 2018). (D) Scatter plots showing the correlation of the fold enrichment values against the input signals (ChIP/Input) (TSS±2 kb) of H2AK119ub1 between control and *Pcgf1^Δ/Δ^* LSK cells. PRC1 targets are indicated in a green box. (E) Snapshots of Flag-PCGF1 and H2AK119ub1 ChIP signals at the *Cebpa* and *Cebpe* gene loci. (F) ChIP qPCR assays for H2AK119ub1 at the *Cebpa* promoter in control and *Pcgf1^Δ/Δ^* LSK cells. The relative amounts of immunoprecipitated DNA are depicted as a percentage of input DNA. Data are shown as the mean ± SEM (n=3). (G) Box-and-whisker plots showing ATAC signal levels at proximal promoters (TSS ± 2kb) in *Pcgf1^Δ/Δ^* CD135^-^LSK cells relative to those in control cells. The data of all ATAC peaks (n=18,417), PCGF1 target genes (TG) (n=670), and non-PCGF1 TG (n=17,747) are shown. Boxes represent 25-75 percentile ranges. The whiskers represent 10-90 percentile ranges. Horizontal bars represent medians. Mean values are indicated by red crosses. (H) Top DNA motifs identified in ATAC peaks at proximal promoters (TSS ± 2kb) positively or negatively enriched in *Pcgf1^Δ/Δ^* CD135^-^LSK cells compared to corresponding controls. (I) The ATAC signal levels of peacks containing CEBP binding motif (GSE21512, n=233) in CD135^-^LSK cells from control and *Pcgf1^Δ/Δ^*mice. \**p*<0.05; \*\**p*<0.01; \*\*\**p*<0.001; n.s., not significant by the Student’s *t*-test.

We then defined “PCGF1 targets” whose H2AK119ub1 levels were decreased by *Pcgf1* deletion greater than twofold at promoter regions in HSPCs (997 RefSeq ID genes; Table S2). We found that *Cebpa* and *Cebpe* were included in PCGF1 targets, showed reductions in H2AK119ub1 levels, and were de-repressed in expression in *Pcgf1^Δ/Δ^* LSK cells (Fig 3D and E, 2C and S5C). ChIP-qPCR confirmed a significant reduction of H2AK119ub1 levels at the promoter region of *Cebpa* in *Pcgf1^Δ/Δ^* HSPCs (Fig 3F). These results indicate that the deletion of *Pcgf1* compromises PRC1.1 function and causes precocious activation of key myeloid regulator genes in HSPCs.

To clarify whether the *Pcgf1* deletion has any impact on the chromatin accessibility in HSPCs, we performed an assay for transposase accessible chromatin with high-throughput sequencing (ATAC-seq) analysis using CD135^−^LSK HSPCs, which include HSCs and MPP1-3 but not lymphoid-primed MPP4. ATAC-seq profiles open chromatin regions enriched for transcriptional regulatory regions, such as promoters and enhancers. ATAC peaks were significantly enriched at the promoter regions of PCGF1 target genes, but not of PCGF1 non-target genes, in *Pcgf1^Δ/Δ^* HSPCs compared to the corresponding controls (Fig 3D), further validating derepression of PCGF1 targets upon the deletion of *Pcgf1*. Motif analysis of ATAC peaks in the proximal promoter regions (TSS±2 kb) revealed that the CTCF motif, which has been reported to be associated with differentiation of HSCs (Buenrostro et al., 2018; Takayama et al., 2021), was highly enriched in *Pcgf1^Δ/Δ^*HSPCs (Fig 3H). Interestingly, the other top-ranked motifs were related to stress response transcription factors, such as Bach family (Bach1 and 2), CNC family (Nrf2 and NF-E2), AP1 family, and Atf3 (Fig 3H and Table S3). In contrast, the binding motifs for transcription factors essential for T and B-cell commitment, including HEB and E2A (de Pooter & Kee, 2010), were negatively enriched in *Pcgf1^Δ/Δ^* HSPCs (Fig 3H). Moreover, ATAC-seq analyses also revealed trend toward enrichment of ATAC peaks containing CEBP motif in *Pcgf1^Δ/Δ^* HSPCs compared to the control, suggesting that chromatin regions containing CEBP motif tended to open in *Pcgf1^Δ/Δ^*HSPCs (Fig 3I). Together with the significant up-regulation of *Cebpa* and *Cebpe* in *Pcgf1^Δ/Δ^*HSPCs (Figs. 2C and S5C), these results further support the notion that PRC1.1 deficiency caused precocious activation of myeloid differentiation program at the expense of lymphoid differentiation program in HSPCs.

### PCGF1 inhibition facilitates emergency myelopoiesis

Myeloid-biased hematopoiesis in *Pcgf1*-deficient mice reminded us of the myeloproliferative reactions caused by emergencies such as regeneration (Manz & Boettcher, 2014). Individual GMPs scatter throughout the BM in the steady state, while expanding GMPs evolve into GMP clusters during regeneration, which, in turn, differentiate into granulocytes. Inducible activation of β-catenin and *Irf8* controls the formation and differentiation of GMP clusters, respectively (Hérault et al., 2017). Immunofluorescence analyses of BM sections readily detected GMP clusters in steady-state *Pcgf1^Δ/Δ^* BM (Fig 4A), which was reminiscent of those observed during regeneration after 5-fluorouracil (5-FU) treatment (Fig S6A). To address if PCGF1 also regulates myelopoiesis at the GMP level during regeneration, we challenged control and *Pcgf1^Δ/Δ^*mice with a single dose of 5-FU (Fig 4B). Consistent with the previous report (Hérault et al., 2017), control mice showed transient expansion of BM HSPCs and GMPs around day 14 and subsequent burst of circulating PB myeloid cells around day 21. In sharp contrast, *Pcgf1^Δ/Δ^* mice displayed sustained GMP expansion until day 28 without efficient production of PB myeloid cells, leading to the accumulation of excess GMPs (Fig 4C).

**Figure 4.**
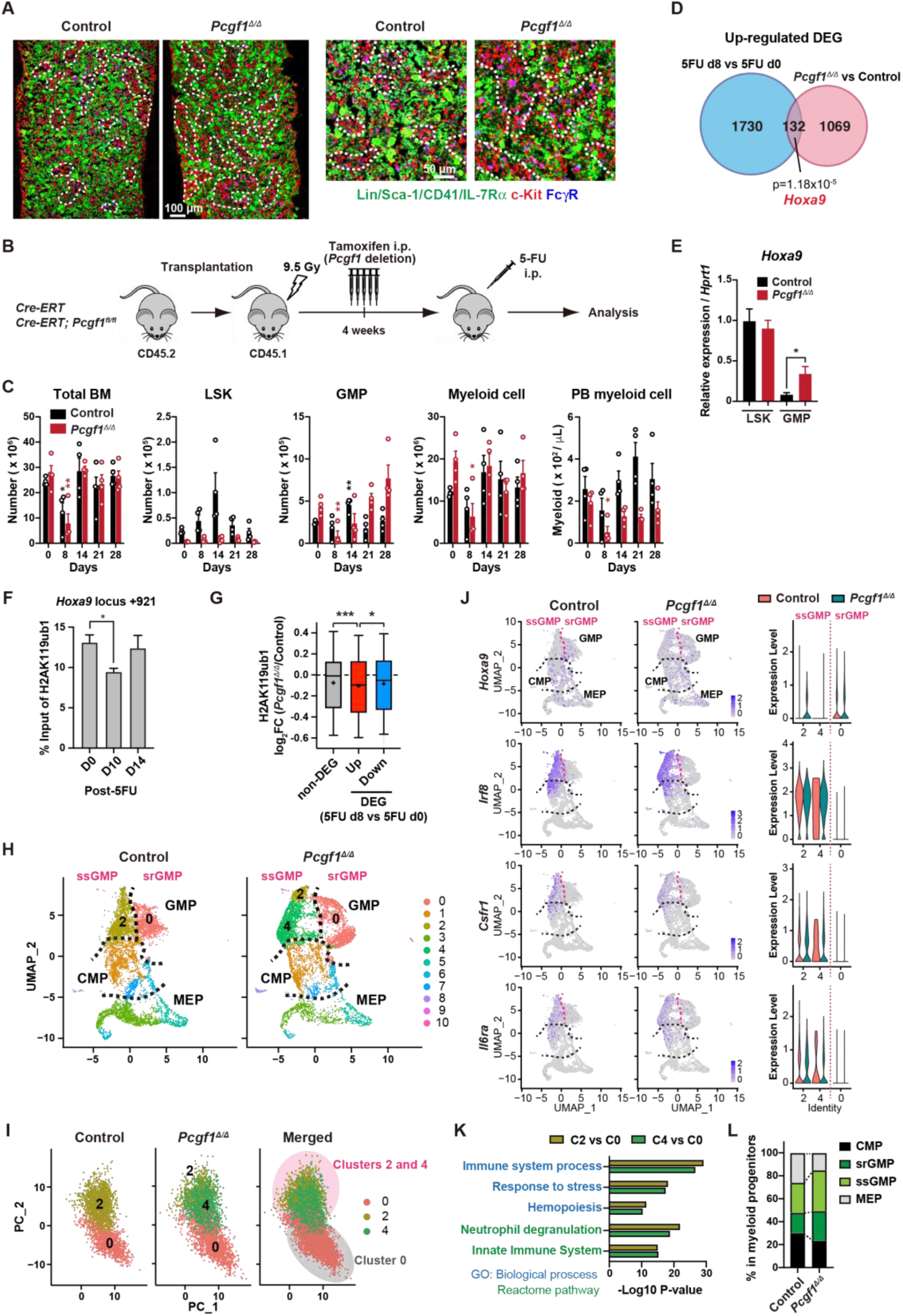
PCGF1 negatively regulates GMP self-renewal. (A) Immunofluorescence staining of BM sections from control and *Pcgf1^Δ/Δ^* mice. Magnified images are depicted in the right panels. Dotted lines denote clusters of GMP (Lin^−^Sca-1^−^CD41^−^IL-7Rα^−^c-Kit^+^FcγR^+^) (c-Kit, red; FcγR, blue; merged, purple). (B) Strategy to analyze emergency myelopoiesis induced by one shot of 5-FU (150 mg/kg). (C) Absolute numbers of total BM cells, LSK cells, GMPs, and Mac-1^+^ myeloid cells in a unilateral pair of femur and tibia and Mac-1^+^ myeloid cells in PB at the indicated time points post-5-FU injection). Data are shown as the mean ± SEM (n=4). Each symbol is derived from an individual culture. (D) A venn diagram showing the overlap between up-regulated DEGs in 5-FU treated GMPs on day 8 and up-regulated DEGs in *Pcgf1^Δ/Δ^* GMPs. (E) Quantitative RT-PCR analysis of *Hoxa9* in LSK cells and GMPs. *Hprt1* was used to normalize the amount of input RNA. Data are shown as the mean ± SEM (n=3). (F) ChIP qPCR assays for H2AK119ub1 at the *Hoxa9* locus in GMPs from WT mice on days 0, 10, and 14 post-5-FU treatment. The relative amounts of immunoprecipitated DNA are depicted as a percentage of input DNA. Data are shown as the mean ± SEM (n=3). (G) Fold changes in H2AK119ub1 levels in *Pcgf1^Δ/Δ^* GMPs relative to control GMPs at the promoters of non-DEGs and DEGs up- and down-regulated in day 8 GMPs compared to day 0 GMPs post-5-FU treatment. (H) UMAP plots illustrating the identification of cell clusters based on single cell transcriptomic profiling of control and *Pcgf1^Δ/Δ^* myeloid progenitors (Lin^−^Sca-1^−^c-Kit^+^). (I) PCA plots of control and *Pcgf1^Δ/Δ^* GMPs individually and in combination. (J) UMAP and violin plots showing expression of *Hoxa9, Irf8, Csf1r*, and *Il-6ra* in control and *Pcgf1^Δ/Δ^* myeloid progenitors. (K) Gene Ontology and pathway enrichment analyses using DEGs in the indicated clusters. (L) Proportion of CMPs, MEPs, ssGMPs and srGMPs in control and *Pcgf1^Δ/Δ^* myeloid progenitors. \**p*<0.05; \*\**p*<0.01; \*\*\**p*<0.001 by the Student’s *t*-test.

GMPs can be divided into steady-state GMPs (ssGMP) and self-renewing GMPs (srGMP), the latter of which transiently increase during regeneration (Hérault et al., 2017). We performed RNA-seq analysis of GMPs isolated from 5-FU-treated WT mice at various time points (Fig S6B and C), and defined differentially expressed genes (DEGs) on day 8 after 5-FU treatment, since srGMPs reportedly most expand on that day (Fig S6B and C; and Table S4) (Hérault et al., 2017). Of note, a significant portion of upregulated DEGs in day 8 5-FU-treated GMPs were also upregulated in *Pcgf1^Δ/Δ^*GMPs (Fig 4D and Table S4). These overlapping genes included *Hoxa9*, a PRC1.1 target (Fig S6D) (Ross et al., 2012; Shinoda et al., 2022; Tara et al., 2018), which is highly expressed in srGMPs (Hérault et al., 2017). RT-qPCR confirmed significantly higher expression of *Hoxa9* in *Pcgf1^Δ/Δ^* GMPs than control GMPs (Fig 4E). Correspondingly, 5-FU treatment transiently decreased H2AK119ub1 levels at *Hoxa9* locus around day 10 (Fig 4F). ChIP-seq analysis also revealed significant reductions in H2AK119ub1 levels at promoters of upregulated DEGs in day 8 5-FU-treated GMPs in *Pcgf1^Δ/Δ^* GMPs (Fig 4G). These results suggest that transient inhibition of PRC1.1 de-represses genes critical to expand srGMP during myeloid regeneration, although the expression of PRC1.1 genes remained largely unchanged during regeneration (Fig S6E).

To better understand the role of PRC1.1 in myeloid progenitors, we performed single cell RNA-seq (scRNA-seq) of Lin^−^Sca-1^−^c-Kit^+^ myeloid progenitors from control and *Pcgf1^Δ/Δ^* mice at steady state. We used data from 6,171 control and 6,198 *Pcgf1^Δ/Δ^* single cells and identified 10 major clusters based on dimension reduction by UMAP (Fig 4H). Functional annotation of respective UMAP clusters using previously reported myeloid progenitor cell gene expression profiles (Nestorowa et al., 2016) assigned clusters 0, 2 and 4 to GMPs (Fig 4H). PCA analysis subdivided GMPs into two major groups (Fig 4I). These two groups exhibited distinct expression profiles of *Hoxa9*, *Irf8*, *Csf1r*, and *Il6ra*, key genes differentially expressed between ssGMPs (*Hoxa9*^lo^, *Irf8*^hi^, *Csf1r*^hi^, *Il6ra*^hi^) and srGMPs (*Hoxa9*^hi^, *Irf8*^lo^, *Csf1r*^lo^, *Il6ra*^lo^) (Hérault et al., 2017), and we classified clusters 2 and 4 as ssGMPs and cluster 0 as srGMPs (Fig 4J). Gene Ontology and pathway enrichment analyses using DEGs (cluster 2 or 4 versus cluster 0) revealed that clusters 2 and 4 represented more mature myeloid cell populations than cluster 0 (Fig 4K). As expected, *Pcgf1^Δ/Δ^* myeloid progenitors had a greater proportion of total GMPs including srGMPs than controls (Fig 4L). Of note, the frequency of srGMPs was also increased in *Pcgf1^Δ/Δ^* myeloid progenitors (Fig 4L). These results indicate that PRC1.1 restricts expansion of self-renewing GMPs and suggest that transient PRC1.1 inhibition allows for temporal amplification of GMPs and their subsequent differentiation to mature myeloid cells.

### PCGF1 restricts GMP self-renewal network

To further investigate the mechanism by which PCGF1 regulates GMPs, we took advantage of *in vitro* culture experiments. Remarkably, while control GMPs stopped proliferation on day 4, *Pcgf1^Δ/Δ^* GMPs kept growing until day 11 (Fig 5A). Moreover, while comparable numbers of GMPs (CD34^+^FcγR^+^c-Kit^+^Sca-1^-^Lineage^-^) were produced by control and *Pcgf1^Δ/Δ^* HSCs on day 7 of culture, control HSCs showed a rapid decline in GMP production afterward but *Pcgf1^Δ/Δ^*HSCs persistently produced GMPs until day 23 (Fig 5B). Of interest, differentiation of the expanded *Pcgf1^Δ/Δ^* GMPs was largely blocked, as indicated by reduced Mac1^+^ differentiated myeloid cells/GMP ratios until day 19 (Fig S6F), suggesting *Pcgf1^Δ/Δ^*GMPs underwent enhanced self-renewal rather than differentiation. Furthermore, *Pcgf1^Δ/Δ^* HSPCs showed remarkably sustained colony formation activity upon serial replating with myeloid cytokines, which is in line with the elevated self-renewing activity of *Pcgf1^Δ/Δ^* GMPs (Fig 5C).

**Figure 5.**
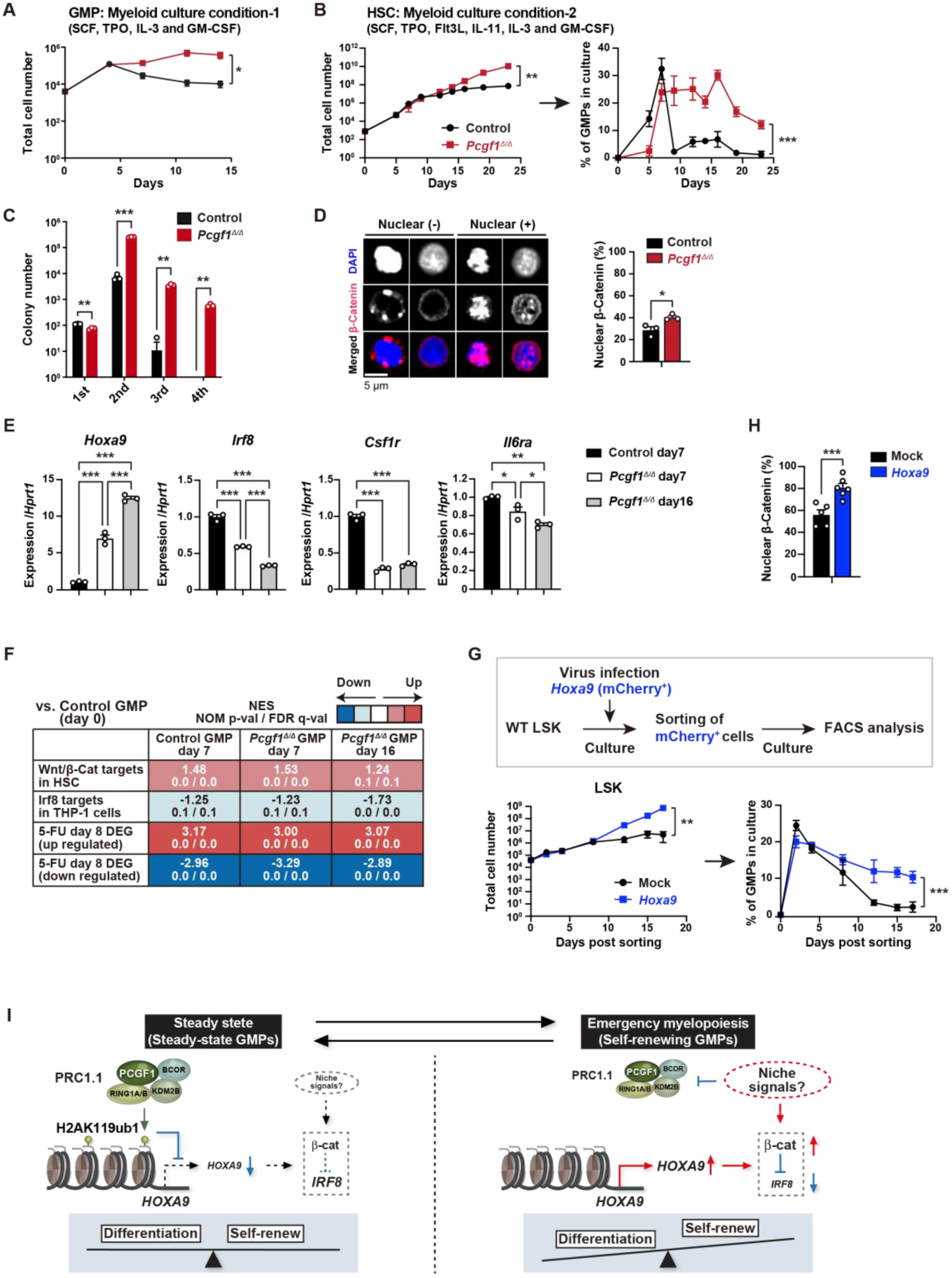
PCGF1 restricts self-renewal of GMPs by attenuating *Hoxa9* expression. (A) Growth of control and *Pcgf1^Δ/Δ^* GMPs in culture. Cells were cultured in triplicate under myeloid culture condition-1 (20 ng/mL SCF, TPO, IL-3, and GM-CSF). Data are shown as the mean ± SEM. Growth of control and *Pcgf1^Δ/Δ^* HSCs under myeloid culture condition-2 (25 ng/mL SCF, TPO, Flt3L, and IL-11 and 10 ng/mL IL-3 and GM-CSF). Cells were cultured in triplicate. The proportion of GMPs in culture is depicted on the right panel. (C) Replating assay data. Seven hundred LSK cells were plated in a methylcellulose medium containing 20 ng/mL of SCF, TPO, IL-3, and GM-CSF. After 10 days of culture, colonies were counted and pooled, and 1×10^4^ cells were then replated in the same medium every 7 days. (D) Proportion of GMPs with nuclear β-catenin in control and *Pcgf1^Δ/Δ^* GMPs in HSC culture on day16 in (B). Representative immunofluorescent signals of β-catenin are shown on the right panel. (E) Quantitative RT-PCR analysis of *Hoxa9, Irf8, Csf1r*, and *Il-6ra* in control and *Pcgf1^Δ/Δ^* GMPs in HSC culture in (B) at the indicated time points. *Hprt1* was used to normalize the amount of input RNA. Data are shown as the mean ± SEM (n=3). (F) GSEA using RNA-seq data. The gene sets used are indicated in Supplementary Table 1. (G) Growth of mock control and *Hoxa9*-expressing LSK cells. LSK cells transduced with a *Hoxa9* retrovirus harboring mCherry marker gene were cultured in triplicate under myeloid culture condition-2. The proportion of GMPs in culture is depicted on the right panel. (H) Proportion of GMPs with nuclear β-catenin in mock control and *Hoxa9*-expressing GMPs in LSK culture on day 12 in (G). (I) Model of the molecular network controlling GMP self-renewal and differentiation. \**p*<0.05; \*\**p*<0.01; \*\*\**p*<0.001 by the Student’s *t*-test (A-D, H, and G) or the One-way ANOVA (E). Each symbol is derived from an individual culture.

srGMPs have increased levels of nuclear β-catenin, which is known to confer aberrant self-renewal features to leukemic GMPs (Wang et al., 2010) and directly suppresses *Irf8* expression (Hérault et al., 2017). The proportion of nuclear β-catenin was significantly increased in *Pcgf1^Δ/Δ^* GMPs at later time points of culture when GMPs in control culture shrunk but GMPs in *Pcgf1^Δ/Δ^* culture kept expanding (Fig 5B and D). *Pcgf1^Δ/Δ^* GMPs in culture possessed a transcriptional profile typical to srGMPs; up-regulation of *Hoxa9* and down-regulation of *Irf8*, *Csf1r*, and *Il6ra* (Fig 5E). GSEA revealed activation of the Wnt/β-catenin pathway (Shooshtarizadeh et al., 2019) and down-regulation of the Irf8 targets (Kubosaki et al., 2010) in *Pcgf1^Δ/Δ^* GMPs (Fig 5F). We hypothesized that *Hoxa9*, a direct target of PCGF1, could have a role in the GMP self-renewal network. Overexpression of *Hoxa9* in HSPCs significantly enhanced their growth and induced persistent production of GMPs for a long period (Fig 5G and S6G). Most *Hoxa9*-overexpressing GMPs had nuclear β-catenin (Fig 5H). These results indicate that HOXA9 can reinforce activation of β-catenin, thus placing HOXA9 as a component of the GMP self-renewal network and PCGF1-PRC1.1 as a negative regulator of this network (Fig 5 I).

### Constitutive PCGF1 loss promotes malignant transformation

In leukemia, GMP clusters are constantly produced owing to persistent activation of the myeloid self-renewal network and a lack of termination cytokines that normally restore HSC quiescence (Hérault et al., 2017). A significant portion of *Pcgf1^Δ/Δ^* mice, which exhibit constant production of GMP clusters, developed lethal Myeloproliferative neoplasms (MPN) with severe anemia and massive accumulation of mature myeloid cells in PB, BM, and spleen (Fig 6A-E). Accumulation of myeloid cells was evident in BM and spleen sections (Fig 6F). A part of *Pcgf1^Δ/Δ^* mice also developed lethal T-cell acute lymphoblastic leukemia (T-ALL) (Fig 6A and G) like *Bcor* mutant mice (Tara et al., 2018). These results indicate that constitutive activation of the GMP self-renewal network in *Pcgf1^Δ/Δ^* mice serves to promote malignant transformation. Taken together, these results highlight the importance of PRC1.1-dependent suppression of the myeloid self-renewing network to prevent malignant transformation.

**Figure 6.**
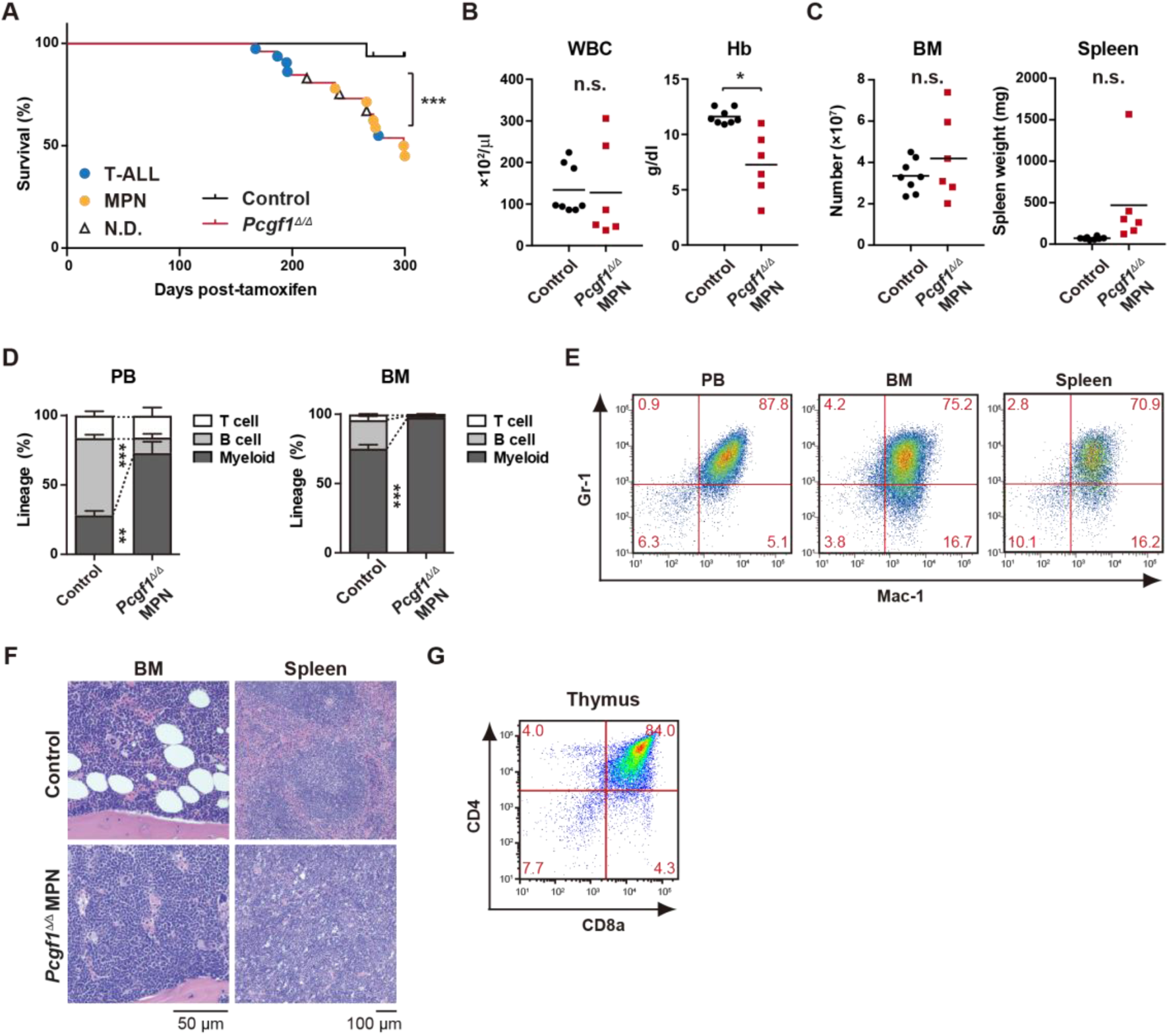
Development of lethal myeloproliferative neoplasm in *Pcgf1^Δ/Δ^* mice. (A) Kaplan-Meier survival curves of control (n=7) and *Pcgf1^Δ/Δ^*(n=25) mice after the tamoxifen injection. (B) White blood cell (WBC) and hemoglobin (Hb) in PB from control (n=8) and moribund *Pcgf1^Δ/Δ^* MPN mice (n=6). Bars indicate median values. Data are shown as the mean ± SEM. (C) Absolute numbers of total BM cells and spleen weight in control (n=8) and moribund *Pcgf1^Δ/Δ^* MPN mice (n=6). (D) The proportions of Mac-1^+^ and/or Gr-1^+^ myeloid cells, B220^+^ B cells, and CD4^+^ or CD8^+^ T cells in PB and BM in control (n=8) and moribund *Pcgf1^Δ/Δ^* MPN mice (n=6). (E) Representative flow cytometric profiles of PB, BM, and spleen of control and moribund *Pcgf1^Δ/Δ^* MPN mice. The percentages of gated populations over CD45.2^+^ live cells are indicated. (F) Representative histology of BM and spleen from control and moribund *Pcgf1^Δ/Δ^* MPN mice observed by hematoxylin-eosin staining. (G) Representative flow cytometric profiles of thymus from control mice and moribund *Pcgf1^Δ/Δ^* T-ALL mice. \**p*<0.05; \*\**p*<0.01; \*\*\**p*<0.001 by the Student’s *t*-test. Each symbol is derived from an individual mouse.

## DISCUSSION

In this study, we demonstrated that PCGF1 contributes to balanced hematopoiesis by restricting precocious myeloid commitment of HSPCs and expansion of myeloid progenitors while its inhibition promotes emergency myelopoiesis and myeloid transformation. These findings present a sharp contrast with PCGF4/BMI1 essential for self-renewal of HSCs (Iwama et al., 2004; Oguro et al., 2006; Park et al., 2003) and underscore distinct functions between canonical PRC1 and non-canonical PRC1.1 in hematopoiesis (Fig S6H).

PcG and trithorax group proteins mark developmental regulator gene promoters with bivalent histone domains to keep them poised for activation in ES cells(Bernstein et al., 2006). We previously reported that canonical PRC1 reinforces bivalent domains at the B cell regulator genes, *Ebf1* and *Pax5*, to maintain B-cell lineage commitment poised for activation in HSPCs (Oguro et al., 2010). In contrast, PCGF1 appeared to target non-bivalent PRC1 target genes marked with moderate levels of H2AK119ub1 and H3K27me3. Among these, PCGF1 targets myeloid regulator genes, such as *Cebpa*, thereby negatively regulating myeloid commitment. Our findings indicate that canonical and non-canonical PRC1 restrict the lymphoid and myeloid commitment of HSPCs, respectively, by targeting different transcriptional programs of differentiation, thereby fine-tuning the balance of HSPC commitment (Fig S6H). Although there might be considerable functional redundancy between canonical and non-canonical PRC1 complexes, our results uncovered a unique function of PRC1.1 in the lineage commitment of HSPCs.

Myeloid-biased output from HSPCs is one of the hallmarks of emergency hematopoiesis (Trumpp et al., 2010; Zhao & Baltimore, 2015). In mouse models of regeneration, myeloid-biased MPP2 and MPP3 are transiently overproduced, suggesting that HSCs produce functionally distinct lineage-biased MPPs to adapt blood production to hematopoietic demands (Pietras et al., 2015). In the present study, we found that *Pcgf1*-deficient hematopoiesis recapitulates sustained emergency myelopoiesis, although the production of circulating myeloid cells was not enhanced. Expanding GMPs, GMP clusters during regeneration, which, in turn, differentiate into granulocytes (Hérault et al., 2017). Of note, PCGF1 loss induced constitutive GMP cluster formation at steady state and sustained GMP expansion in mice after myeloablation and in culture. Correspondingly, *Pcgf1*-deficient mice had a greater number of self-renewing GMPs than control mice. This unique phenotype may implicate the importance of transient but not constitutive PCGF1 repression for proper myeloid regeneration. β-catenin and *Irf8* constitute an inducible self-renewal progenitor network controlling GMP cluster formation, with β-catenin directly suppressing *Irf8* expression while restoration of *Irf8* expression terminating the self-renewal network and inducing GMP differentiation (Hérault et al., 2017) (Fig 5I). *Hoxa9*, which is up-regulated in srGMPs, is one of the PRC1.1 targets in myeloid progenitors (Fig S6) (Ross et al., 2012; Shinoda et al., 2022; Tara et al., 2018). We demonstrated that *Hoxa9* expression activates β-catenin and promotes GMP self-renewal, identifying HOXA9 as a component of the GMP self-renewal network. Of note, PRC1.1 is transiently inhibited to de-repress such GMP self-renewal network genes. This transient nature of PRC1.1 inhibition allows for srGMP expansion and GMP cluster formation followed by proper differentiation of expanded GMPs. As expression levels of PRC1.1 components remained unchanged during hematopoietic regeneration, non-canonical PRC1.1 activity could be modulated by posttranslational modifications in response to extracellular stimuli like canonical PRC1 (Banerjee Mustafi et al., 2017; Liu et al., 2012; Nacerddine et al., 2012; Voncken et al., 2005). How extrinsic signals modulate PRC1.1 functions to regulate myelopoiesis remains an important question.

The molecular machineries that drive emergency myelopoiesis are often hijacked by transformed cells (Hérault et al., 2017). A significant portion of *Pcgf1*-deficient mice eventually developed lethal MPN after a sustained myeloproliferative state. These findings indicate that PRC1.1 functions as a critical negative regulator of myeloid transformation. Among the components of PRC1.1, *BCOR* and *BCLRL1*, but not *PCGF1* are targeted by somatic gene mutations in various hematological malignancies, including myelodysplastic syndrome (MDS), chronic myelomonocytic leukemia (CMML), and acute myeloid leukemia (AML) (Isshiki & Iwama, 2018). We reported that mice expressing a carboxyl-terminal truncated BCOR, which cannot interact with PCGF1, showed myeloid-biased hematopoiesis like *Pcgf1*-deficient mice. Importantly, HSPCs in these mice showed a growth advantage in the myeloid compartment, which was further enhanced by the concurrent deletion of *Tet2*, leading to the development of lethal MDS (Tara et al., 2018). De-repression of myeloid regulator genes, such as *Cebp* family and *Hoxa* cluster genes, were also detected in *Bcor* mutant progenitor cells (Tara et al., 2018). These findings also support the idea that PRC1.1 restricts myeloid transformation by transcriptionally repressing aberrant activation of myeloid regeneration programs.

Collectively, our findings highlight a critical role of PRC1.1 in coordinating steady-state and emergency hematopoiesis and preventing malignant transformation. They also suggest that transient inhibition of PRC1.1 would be a novel approach to temporarily induce emergency myelopoiesis and enhance myeloid cell supply while avoiding the potential risk for malignant transformation.

## MATERIALS AND METHODS

### Mice

Wild-type mice (C57BL/6) and *Rosa::Cre-ERT2* mice were purchased from the Japan SLC and TaconicArtemis GmbH, respectively. *Pcgf1^fl^*and *Cebpa^fl^* mice were kindly provided by Haruhiko Koseki and Daniel G. Tenen, respectively, and previously reported (Almeida et al., 2017; Zhang et al., 2004). All experiments using mice were performed in accordance with our institutional guidelines for the use of laboratory animals and approved by the Review Board for Animal Experiments of Chiba University (approval ID: 30-56) and the University of Tokyo (approval ID: PA18-03).

### Bone marrow transplantation

To generate hematopoietic cell-specific *Pcgf1* KO mice, we transplanted total BM cells (5×10^6^) from *Rosa::Cre-ERT* and *Cre-ERT;Pcgf1^fl/fl^* mice into lethally irradiated (9.5 Gy) CD45.1 recipient mice. For competitive bone marrow transplantation assay, we transplanted total BM cells (2 x 10^6^) from CD45.2 donor mice with CD45.1^+^ competitor total BM cells (2×10^6^) into lethally irradiated (9.5 Gy) CD45.1 recipient mice. To induce Cre activity, transplanted-mice were injected with 100 μL of tamoxifen (Sigma-Aldrich) dissolved in corn oil (Sigma-Aldrich) at a concentration of 10 mg/mL intraperitoneally once a day for 5 consecutive days 4 weeks after transplantation.

### Locus-specific genotyping of *Pcgf1, Cebpa* and *Rosa::Cre-ERT*

To detect *Pcgf1^fl^*, *Pcgf1^Δ^*, *Cebpa^fl^*, *Cebpa^Δ^* and *Rosa::Cre-ERT* PCR reactions were performed using the specific oligonucleotides. The oligonucleotide sequences used were shown in Table S5.

### 5-FU challenge

8-12-week old wild-type mice or control and *Pcgf1^Δ/Δ^*were injected with 300 μL PBS or 150 mg/kg (3.75 mg per 25 g body weight mouse) 5-FU (Kyowa KIRIN) dissolved in 300 μL PBS intraperitoneally once.

### Flow cytometry analyses and antibodies

The monoclonal antibodies recognizing the following antigens were used in flow cytometry and cell sorting: CD45.1(A20), CD45.2 (104), Gr-1 (RB6-8C5), CD11b/Mac-1 (M1/70), Ter-119 (TER-119), B220 (RA3-6B2), CD127/IL-7R (SB/119), CD4 (GK1.5), CD8a (53–6.7), CD117/c-Kit (2B8), Sca-1 (D7), CD34 (RAM34), CD150 (TC15-12F12.2), CD48 (HM48-1), CD135 (A2F10), CD16/32/FcγRII-III (93), CD41 (eBioMWReg30), CD105 (MJ7/18), Ly6D (49-H4), lineage mixture (Gr-1, Mac-1, Ter-119, CD127/IL-7R, B220, CD4, CD8α) and lineage mixture for CLP (Gr-1, Mac-1, Ter-119, B220, CD4, CD8α). Monoclonal antibodies were purchased from BioLegend, Tonbo Biosciences, Thermo Fisher Scientific or BD Bioscience. Dead cells were eliminated by staining with 0.5 μg/mL Propidium iodide (Sigma-Aldrich). All flow cytometric analyses and cell sorting were performed on FACSAria IIIu, FACSCanto II and FACSCelesta (BD Bioscience). Cell surface protein expression used to define hematopoietic cell types were as follows:

HSC: CD150^+^CD48^−^CD135^−^CD34^−^c-Kit^+^Sca-1^+^Lineage^−^
MPP1: CD150^+^CD48^−^CD135^−^CD34^+^c-Kit^+^Sca-1^+^Lineage^−^
MPP2: CD150^+^CD48^+^CD135^−^CD34^+^c-Kit^+^Sca-1^+^Lineage^−^
MPP3: CD150^−^CD48^+^CD135^−^CD34^+^c-Kit^+^Sca-1^+^Lineage^−^
MPP4: CD150^−^CD48^+^CD135^+^CD34^+^c-Kit^+^Sca-1^+^Lineage^−^
CMP: CD34^+^FcγR^−^c-Kit^+^Sca-1^−^Lineage^−^
GMP: CD34^+^FcγR^+^c-Kit^+^Sca-1^−^Lineage^−^
MEP: CD34^−^FcγR^−^c-Kit^+^Sca-1^−^Lineage^−^
pre-GM: CD150^−^CD105^−^FcγR^−^CD41^−^c-Kit^+^Sca-1^−^Lineage^−^
CLP: c-Kit^low^Sca-1^low^CD135^+^IL7R^+^Lineage (for CLP)^−^
ALP: Ly6D^−^cKit^low^Sca-1^low^CD135^+^IL7R^+^Lineage^−^
BLP: Ly6D^+^cKit^low^Sca-1^low^CD135^+^IL7R^+^Lineage^−^
LSK: c-Kit^+^Sca-1^+^Lineage^−^
LK: c-Kit^+^Lineage^−^
Pro-B: B220^+^CD43^+^IgM^−^
Pre-B: B220^+^CD43^−^IgM^−^

### Quantitative RT-PCR

Total RNA was extracted using a RNeasy Micro Plus Kit (QIAGEN) or TRIZOL LS solution (MOR) and reverse transcribed by the SuperScript IV First-Strand Synthesis System (Invitrogen) or the ReverTra Ace α-(TOYOBO) with an oligo-dT primer. Real-time quantitative PCR was performed with a StepOnePlus Real-Time PCR System (Life Technologies) using FastStart Universal Probe Master (Roche) and the indicated combinations of the Universal Probe Library (Roche), or TB Green Premix Ex Taq II (TaKaRa Bio). All data are presented as relative expression levels normalized to *Hprt* expression. The primer sequences used were shown in Table S5 (Murakami et al., 2021; Sonntag et al., 2018).

### Limiting dilution assay

For *in vivo* limiting dilution assay, we transplanted limiting numbers of total BM cells (1×10^4^, 4×10^4^, 8×10^4^, and 2×10^5^) isolated from primary recipients (control and *Pcgf1^Δ/Δ^*mice 1 month after tamoxifen injections) with CD45.1^+^ competitor total BM cells (2×10^5^) into lethally irradiated CD45.1 recipient mice. PB analyses were performed at 16 weeks after transplantation.

For *in vitro* limiting dilution assay, we sorted HSCs from control and *Pcgf1^Δ/Δ^* mice 1 month after tamoxifen injections and cultured limiting numbers of the cells (1, 5, 25, and 125) with TSt-4 (B cells) or TSt-4/DLL1 stromal cells (T cells) in RPMI (Thermo Fisher Scientific) supplemented with 10% BSA (093001; STEMCELL Technologies), 50 μM 2-ME (Sigma-Aldrich), 100 μM MEM Non-Essential Amino Acids solution (Gibco), 100 μM sodium pyruvate (Gibco) and 2 ng/mL recombinant mouse IL-7 (577802; BioLegend) for 28 days. The generation of CD19^+^ B cells or Thy1.2^+^ T cells in each well was detected by flow cytometry.

### Cell cycle assay

BM cells were stained with antibodies against cell-surface markers. After washing, cells were fixed and permeabilized with a BD Phosflow Lyse/Fix Buffer and a BD Phosflow Perm Buffer II (BD Bioscience) according to the manufacturer’s instructions. Cells were stained with FITC-Ki67 antibody (#11-5698-82; Thermo Fisher Scientific) at room temperature for 30 min and then with 1 μg/mL 7-AAD (Sigma-Aldrich). Flow cytometric analyses were performed on FACSAria IIIu (BD Bioscience).

### Cell culture

For growth assays, sorted CD34^−^CD150^+^LSK HSCs, LSK cells and GMPs were cultured in S-Clone SF-O3 (Sanko Junyaku) supplemented with 0.1% BSA (093001; STEMCELL Technologies), 50 μM 2-ME (Sigma-Aldrich) and 1% penicillin/streptomycin/glutamine (Gibco). 20 ng/mL of recombinant mouse SCF (579706; Biolegend) and recombinant human TPO (763706; Biolegend) for HSC culture conditions and 10 ng/mL of SCF, TPO, recombinant mouse IL-3 (575506; Biolegend), and recombinant murine GM-CSF (315-03; PeproTech) for myeloid culture condition-1 were added to cultures. In the case of myeloid culture condition-2, sorted CD150^+^CD48^−^CD135^−^CD34^−^LSK HSCs were cultured in IMDM (Gibco) supplemented with 5% FBS, 50 μM 2-ME (Sigma-Aldrich), 1% penicillin/streptomycin/glutamine (Gibco), 1 mM sodium pyruvate (Gibco) and 0.1 mM MEM Non-Essential Amino Acids solution (Gibco). 25 ng/mL of SCF, TPO, recombinant human Flt3L (300-19; PeproTech) and recombinant murine IL-11 (220-11; PeproTech) and 10 ng/mL of IL-3 and GM-CSF were added to cultures.

For replating assays, LSK cells were plated in methylcellulose medium (Methocult M3234; STEMCELL Technologies) containing 20 ng/mL of SCF, TPO, IL-3, and GM-CSF.

### Retroviral vector and virus production

Full-length *Pcgf1* and *Bmi1* cDNA tagged with a 3×Flag at the N-terminus was subcloned into the retroviral vector pGCDNsam-IRES-EGFP. Full-length *Hoxa9* cDNA was subcloned into the retroviral vector pMYs-IRES-mCherry. A recombinant retrovirus was generated by a 293gpg packaging cell line. The virus in supernatants of 293gpg cells was concentrated by centrifugation at 6,000g for 16 hours.

### Immunofluorescence imaging of bone marrow and spleen sections

Isolated mouse femurs were immediately placed in ice-cold 2% paraformaldehyde solution (PFA/PBS) and fixed under gentle agitation for 16 hours. The samples were then incubated in 15% and 30% sucrose for cryoprotection overnight. Samples were embedded in O.C.T. (Sakura) and frozen in cooled hexane. The 7 μm frozen sections were generated with a cryostat (Cryostar NX70, Thermo Scientific) using Kawamoto’s tape method (Kawamoto, 2003). Sections on slide glasses were blocked with staining buffer (10% normal donkey serum in TBS) and an Avidin/Biotin Blocking Kit (VECTOR), then stained with biotinylated anti-lineage antibody cocktail and anti-c-Kit antibody (#AF 1356; R&D Systems), or anti-FcγR-AlexaFluor 647 (#101314; Biolegend) in staining buffer overnight at 4 ℃. For secondary staining, sections were incubated with streptavidin-AlexaFluor 488 (#S11223; Invitrogen) and donkey anti-goat AlexaFluor 555 (#A21432; Invitrogen) antibody for 3 hours at room temperature. Finally, sections were incubated with 1 μg/mL DAPI/TBS for 10 minutes and mounted with ProLong Glass Antifade Mountant (Thermo Scientific). Images of sections were captured on a confocal microscope (Dragonfly, Andor or A1Rsi, Nikon) and processed using Fiji.

### Immunofluorescence imaging of purified GMPs

GMPs were sorted directly onto glass slides using BD AriaIIIu. The cells were washed three times with PBS for 5 min between each staining step. Cells were fixed with 4% PFA for 15 min, permeabilized with 0.1% Triton X-100 for 10 min, and then blocked with 3% BSA for 1 h at room temperature. The cells were then incubated with rabbit anti-mouse β-catenin (#9582S; Cell Signaling) primary antibody at 4°C overnight. The cells were then stained with anti-rabbit AF488A (#20015; Biotium) secondary antibody for 2 h at room temperature. After staining with 1 μg/mL DAPI/PBS for 5 min, the cells were mounted with ProLong™ Glass Antifade Mountant (ThermoFisher). DragonFly (Andor, 40x objective) was used for image acquisition.

### Bulk RNA-seq and data processing

Total RNAs were extracted from 1,000-5,000 cells using an RNeasy Plus Micro Kit (QIAGEN) and cDNAs were synthesized using a SMART-Seq v4 Ultra Low Input RNA Kit for Sequencing (Clontech) according to the manufacturer’s instructions. The ds-cDNAs were fragmented using S220 or M220 Focused-ultrasonicator (Covaris), then cDNA libraries were generated using a NEBNext Ultra DNA Library Prep Kit (New England BioLabs) according to the manufacturer’s instructions. Sequencing was performed using HiSeq1500 or HiSeq2500 (Illumina) with a single-read sequencing length of 60bp. TopHat2 (version 2.0.13; with default parameters) was used to map the reads to the reference genome (UCSC/mm10) with annotation data from iGenomes (Illumina). Levels of gene expression were quantified using Cuffdiff (Cufflinks version 2.2.1; with default parameters).Significant expression differences were detected edgeR (version 3.14; with default parameters), with raw counts generated from String Tie. The super-computing resource was provided by the Human Genome Center, the Institute of Medical Science, the University of Tokyo (http://sc.hgc.jp/shirokane.html). The enrichment analysis

### Single cell RNA-seq and data processing

Control (1.2×10^4^) and *Pcgf1^Δ/Δ^* (1.2×10^4^) LK cells were collected for single cell RNA-seq. mRNA were isolated and libraries were prepared according to Chromium Next GEM Single Cell 3ʹ Reagent Kits v3.1 (10× Genomics). Raw data files (Base call files) were demultiplexed into fastq files using Cell Ranger with mkfastq command. Then, “cellranger count” command was used for feature counts, barcode counts with reference “refdata-gex-mm10-2020-A”. Filtered_feature_bc_matrix included 6,565 control LK cells and 7,651 Pcgf1 KO LK cells. We subsampled 6,565 cells from 7,651 Pcgf1 KO LK cells to adjust cell numbers between KO and control LK. Subsequent analyses were performed using Seurat 4.1.0. Quality filtering for each feature and cell was conducted based on these criteria (min.cells = 3 & min.features = 200 & nFeature_RNA > 200 & nFeature_RNA < 10000 & percent.mt < 5). After quality filtering, 6,171 control LKs and 6,198 KO LKs were used for further analysis. Feature counts are log-normalized with the function of “NormalizeData”. 2,000 highly variable features are selected for PCA. PC 1-10 components are used for UMAP and graph-based clustering with the functions of FindNeighbors(object, reduction = “pca”, dims = 1:15) and FindClusters(object, resolution = 0.28). Cluster 0, 2 and 4 cells are extracted and re-analyzed with PCA. Differentially expressed genes are selected with the function of “FindMarkers(object, min.pct = 0.25)”.

### Chromatin immunoprecipitation (ChIP) assays and ChIP-sequencing

ChIP assays for histone modifications were performed as described previously (Aoyama et al., 2018) using an anti-H2AK119ub1 (#8240S; Cell Signaling Technology) and an anti-H3K27me3 (#07-449; Millipore). BM LSK cells were fixed with 1% FA at 37°C for 2 min, lysed in ChIP buffer (10 mM Tris-HCl pH 8.0, 200 mM NaCl, 1 mM CaCl_2_, 0.5% NP-40 substitute and cOmplete proteases inhibitor cocktail) and sonicated for 5 sec × 3 times by a Bioruptor (UCD-300; Cosmo Bio). After then, cells were digested with Micrococcal Nuclease at 37°C for 40 min (New England BioLabs) and added 10 mM EDTA to stop the reaction. After the addition of an equal volume of RIPA buffer (50 mM Tris-HCl pH 8.0, 150 mM NaCl, 2 mM EDTA pH 8.0, 1% NP-40 substitute, 0.5% sodium deoxycholate, 0.1% SDS and cOmplete proteases inhibitor cocktail), cells were sonicated again for 5 sec x 10 times by a Bioruptor. After centrifugation, supernatants were immunoprecipitated at 4°C overnight with Dynabeads Sheep anti–Rabbite IgG (Invitrogen) conjugated with each antibody. Immunoprecipitates were washed with ChIP wash buffer (10 mM Tris-HCl pH 8.0, 500 mM NaCl, 1 mM CaCl_2_, 0.5% NP-40 substitute and cOmplete proteases inhibitor cocktail) 4 times and TE buffer (10 mM Tris-HCl pH8.0 and 1 mM EDTA pH8.0) twice. Bound chromatins and 30 μL of input DNA were suspended in 95 μL and 65 μL elution buffer (50 mM Tris-HCl pH 8.0, 10mM EDTA pH8.0, 1% SDS and 250 mM NaCl), respectively. After the addition of 5 μL of 5M NaCl, the solutions were incubated at 65°C for 4 hours, treated with 25 μg/mL RNase A (Sigma-Aldrich) at 37°C for 30 min and 0.1 mg/mL proteinase K (Roche) at 50 °C for 1 hour and were purified with a MinElute PCR Purification Kit (QIAGEN).

In 3×Flag-Pcgf1ChIP assay, BM LK cells were fixed with 1% FA at 25°C for 10 min, lysed in RIPA buffer and sonicated for 11 sec × 15 times by a homogenizer (NR-50M; Micro-tec Co.). After centrifugation, supernatants were immunoprecipitated at 4°C overnight with Dynabeads Sheep anti–Mouse IgG (Invitrogen) conjugated with an anti-FLAG antibody (Sigma-Aldrich). After that, the samples were treated in the same way as ChIP assays for histone modifications.

In ChIP-qPCR assay, quantitative real-time PCR was performed with a StepOnePlus Thermal Cycler (Thermo Fisher Scientific) using SYBR Premix Ex Taq II or TB Green Premix Ex Taq II (Takara Bio). The primer sequences used were shown in Table S5 (Tara et al., 2018).

ChIP-seq libraries were prepared using a ThruPLEX DNA-seq Kit (Clontech) according to the manufacturer’s instructions. Bowtie2 (version 2.2.6; with default parameters) was used to map the reads to the reference genome (UCSC/mm10). The RPM (reads per million mapped reads) values of the sequenced reads were calculated every 1,000 bp bin with a shifting size of 100 bp using bedtools. In order to visualize with Integrative Genomics Viewer (IGV) (http://www.broadinstitute.org/igv), the RPM values of the immunoprecipitated samples were normalized by subtracting the RPM values of the input samples in each bin and converted to a bigwig file using wigToBigWig tool. The super-computing resource was provided by the Human Genome Center, the Institute of Medical Science, the University of Tokyo (http://sc.hgc.jp/shirokane.html).

### Assay for Transposase Accessible Chromatin with high-throughput (ATAC)-sequencing

BM CD135^−^LSK cells (1.6-3.0 x 10^4^) and GMPs (3.0 x 10^4^) were lysed in cold lysis buffer (10mM Tris-HCl ph7.4, 10 mM NaCl, 3 mM MgCl_2_, 0.1% IGEPAL CA-630) on ice for 10 min. After centrifugation, nuclei pellets were resuspended with 50 μL of transposase reaction mix (25 μL Tagment DNA buffer (illumine), 2.5 μL Tagment DNA enzyme (illumine) and 22.5 μL water), incubated at 37°C for 35 min and were purified with a MinElute PCR Purification Kit (QIAGEN). After the optimization of PCR cycle number using SYBER Green I Nucleic Acid gel Stain (Takara Bio), transposed fragments were amplified using NEBNext High Fidelity 2× PCR Master mix and index primers, and were purified with a MinElute PCR Purification Kit (QIAGEN). Library DNA was sized selected (240-360 bps) with BluePippin (Sage Scince). Sequencing was performed using HiSeq1500 or HiSeq2500 (Illumina) with a single-read sequencing length of 60bp. Bowtie2 (version 2.2.6; with default parameters) was used to map reads to the reference genome (UCSC/mm10) with annotation data from iGenomes (Illumina). Reads mapped to mitochondria were removed. To ensure even processing, reads were randomly downsampled from each sample to adjust to the smallest read number of samples. MACS (version 2.1.1; with default parameters) was used to call peaks in downsampled reads. The catalogue of all peaks called in any samples was produced by merging all called peaks that overlapped by at least one base pair using bedtools. The MACS bdgcmp function was used to compute the fold enrichment over the background for all populations, and the bedtools map function was used to count fragments in the catalogue in each population. Fragment counts at each site in the catalogue were quantile normalized between samples using the PreprocessCore package in R (3.3.2). We used the Homer package with command annotatePeaks.pl using default parameters to annotate regions with promoter and distal labels and the nearest gene, and with command findMotifsGenome.pl using default parameters to identify enriched motifs, and the catalogue of all called peaks as a background.

### Quantification and Statistical Analysis

Statistical tests were performed using Prism version 9 (Graphpad). The significance of difference was measured by the Student’s *t*-test or One-way *ANOVA*. Data are shown as the mean ± SEM. Significance was taken at values of \**p* less than .05, ** *p* less than .01, and *** *p* less than .001.

### Data and Software Availability

RNA sequence, ChIP sequence and ATAC sequence data were deposited in the DDBJ (accession number DRA008518 and DRA013523).

## AUTHOR CONTRIBUTIONS

Y. Nakajima-Takagi performed the experiments, analyzed results, made the figures, and actively wrote the manuscript. M. Oshima, J. Takano, S. Koide, N. Itokawa, S. Uemura, M. Yamashita, S. Andoh, K. Aoyama, Y. Isshiki, D. Shinida, A. Saraya, F. Arai, K. Yamaguchi, Y. Furukawa, and T. Ikawa assisted with the experiments. H. Koseki generated mice. M. Yamashita aided in interpreting the results and worked on the manuscript. A. Iwama conceived of and directed the project, secured funding, and actively wrote the manuscript.

## ACKNOWLEDGMENTS

We would like to thank Dr. Daniel G. Tenen for providing us with *Cebpa* mutant mice. The super-computing resource was provided by The Human Genome Center, The Institute of Medical Science, The University of Tokyo. This work was supported in part by Grants-in-Aid for Scientific Research (#19H05653, #20K08728) and Scientific Research on Innovative Area “Replication of Non-Genomic Codes” (#19H05746) from Japanese Society for the Promotion of Science (JSPS), Japan, and Moonshot project (#21zf0127003h0001) from AMED, Japan

## REFERENCES

Almeida, M., Pintacuda, G., Masui, O., Koseki, Y., Gdula, M., Cerase, A., Brown, D., Mould, A., Innocent, C., Nakayama, M., Schermelleh, L., Nesterova, T. B., Koseki, H., & Brockdorff, N. (2017). PCGF3/5-PRC1 initiates Polycomb recruitment in X chromosome inactivation. Science, 356(6342), 1081–1084. https://doi.org/10.1126/science.aal2512

Andricovich, J., Kai, Y., Peng, W., Foudi, A., & Tzatsos, A. (2016). Histone demethylase KDM2B regulates lineage commitment in normal and malignant hematopoiesis. J Clin Invest, 126(3), 905–920. https://doi.org/10.1172/JCI84014

Aoyama, K., Oshima, M., Koide, S., Suzuki, E., Mochizuki-Kashio, M., Kato, Y., Tara, S., Shinoda, D., Hiura, N., Nakajima-Takagi, Y., Sashida, G., & Iwama, A. (2018). Ezh1 Targets Bivalent Genes to Maintain Self-Renewing Stem Cells in Ezh2-Insufficient Myelodysplastic Syndrome. iScience, 9, 161–174. https://doi.org/10.1016/j.isci.2018.10.008

Avellino, R., & Delwel, R. (2017). Expression and regulation of C/EBPα in normal myelopoiesis and in malignant transformation. Blood, 129(15), 2083–2091. https://doi.org/10.1182/blood-2016-09-687822

Banerjee Mustafi, S., Chakraborty, P. K., Dwivedi, S. K., Ding, K., Moxley, K. M., Mukherjee, P., & Bhattacharya, R. (2017). BMI1, a new target of CK2α. Mol Cancer, 16(1), 56. https://doi.org/10.1186/s12943-017-0617-8

Bernstein, B. E., Mikkelsen, T. S., Xie, X., Kamal, M., Huebert, D. J., Cuff, J., Fry, B., Meissner, A., Wernig, M., Plath, K., Jaenisch, R., Wagschal, A., Feil, R., Schreiber, S. L., & Lander, E. S. (2006). A bivalent chromatin structure marks key developmental genes in embryonic stem cells. Cell, 125(2), 315–326. https://doi.org/10.1016/j.cell.2006.02.041

Blackledge, N. P., Farcas, A. M., Kondo, T., King, H. W., McGouran, J. F., Hanssen, L. L. P., Ito, S., Cooper, S., Kondo, K., Koseki, Y., Ishikura, T., Long, H. K., Sheahan, T. W., Brockdorff, N., Kessler, B. M., Koseki, H., & Klose, R. J. (2014). Variant PRC1 complex-dependent H2A ubiquitylation drives PRC2 recruitment and polycomb domain formation. Cell, 157(6), 1445–1459. https://doi.org/10.1016/j.cell.2014.05.004

Blackledge, N. P., Rose, N. R., & Klose, R. J. (2015). Targeting Polycomb systems to regulate gene expression: modifications to a complex story. Nat Rev Mol Cell Biol, 16(11), 643–649. https://doi.org/10.1038/nrm4067

Brown, A. L., Wilkinson, C. R., Waterman, S. R., Kok, C. H., Salerno, D. G., Diakiw, S. M., Reynolds, B., Scott, H. S., Tsykin, A., Glonek, G. F., Goodall, G. J., Solomon, P. J., Gonda, T. J., & D’Andrea, R. J. (2006). Genetic regulators of myelopoiesis and leukemic signaling identified by gene profiling and linear modeling. J Leukoc Biol, 80(2), 433–447. https://doi.org/10.1189/jlb.0206112

Buenrostro, J. D., Corces, M. R., Lareau, C. A., Wu, B., Schep, A. N., Aryee, M. J., Majeti, R., Chang, H. Y., & Greenleaf, W. J. (2018). Integrated Single-Cell Analysis Maps the Continuous Regulatory Landscape of Human Hematopoietic Differentiation. Cell, 173(6), 1535–1548.e1516. https://doi.org/10.1016/j.cell.2018.03.074

Busch, K., Klapproth, K., Barile, M., Flossdorf, M., Holland-Letz, T., Schlenner, S. M., Reth, M., Höfer, T., & Rodewald, H. R. (2015). Fundamental properties of unperturbed haematopoiesis from stem cells in vivo. Nature, 518(7540), 542–546. https://doi.org/10.1038/nature14242

Chiba, Y., Mizoguchi, I., Hasegawa, H., Ohashi, M., Orii, N., Nagai, T., Sugahara, M., Miyamoto, Y., Xu, M., Owaki, T., & Yoshimoto, T. (2018). Regulation of myelopoiesis by proinflammatory cytokines in infectious diseases. Cell Mol Life Sci, 75(8), 1363–1376. https://doi.org/10.1007/s00018-017-2724-5

D’Alo’, F., Johansen, L. M., Nelson, E. A., Radomska, H. S., Evans, E. K., Zhang, P., Nerlov, C., & Tenen, D. G. (2003). The amino terminal and E2F interaction domains are critical for C/EBP alpha-mediated induction of granulopoietic development of hematopoietic cells. Blood, 102(9), 3163–3171. https://doi.org/10.1182/blood-2003-02-0479

de Pooter, R. F., & Kee, B. L. (2010). E proteins and the regulation of early lymphocyte development. Immunol Rev, 238(1), 93–109. https://doi.org/10.1111/j.1600-065X.2010.00957.x

Farcas, A. M., Blackledge, N. P., Sudbery, I., Long, H. K., McGouran, J. F., Rose, N. R., Lee, S., Sims, D., Cerase, A., Sheahan, T. W., Koseki, H., Brockdorff, N., Ponting, C. P., Kessler, B. M., & Klose, R. J. (2012). KDM2B links the Polycomb Repressive Complex 1 (PRC1) to recognition of CpG islands. Elife, 1, e00205. https://doi.org/10.7554/eLife.00205

Fukuchi, Y., Shibata, F., Ito, M., Goto-Koshino, Y., Sotomaru, Y., Kitamura, T., & Nakajima, H. (2006). Comprehensive analysis of myeloid lineage conversion using mice expressing an inducible form of C/EBP alpha. EMBO J, 25(14), 3398–3410. https://doi.org/10.1038/sj.emboj.7601199

Gao, Z., Zhang, J., Bonasio, R., Strino, F., Sawai, A., Parisi, F., Kluger, Y., & Reinberg, D. (2012). PCGF homologs, CBX proteins, and RYBP define functionally distinct PRC1 family complexes. Mol Cell, 45(3), 344–356. https://doi.org/10.1016/j.molcel.2012.01.002

Gery, S., Gombart, A. F., Yi, W. S., Koeffler, C., Hofmann, W. K., & Koeffler, H. P. (2005). Transcription profiling of C/EBP targets identifies Per2 as a gene implicated in myeloid leukemia. Blood, 106(8), 2827–2836. https://doi.org/10.1182/blood-2005-01-0358

He, J., Shen, L., Wan, M., Taranova, O., Wu, H., & Zhang, Y. (2013). Kdm2b maintains murine embryonic stem cell status by recruiting PRC1 complex to CpG islands of developmental genes. Nat Cell Biol, 15(4), 373–384. https://doi.org/10.1038/ncb2702

Hérault, A., Binnewies, M., Leong, S., Calero-Nieto, F. J., Zhang, S. Y., Kang, Y. A., Wang, X., Pietras, E. M., Chu, S. H., Barry-Holson, K., Armstrong, S., Göttgens, B., & Passegué, E. (2017). Myeloid progenitor cluster formation drives emergency and leukaemic myelopoiesis. Nature, 544(7648), 53–58. https://doi.org/10.1038/nature21693

Higa, K. C., Goodspeed, A., Chavez, J. S., De Dominici, M., Danis, E., Zaberezhnyy, V., Rabe, J. L., Tenen, D. G., Pietras, E. M., & DeGregori, J. (2021). Chronic interleukin-1 exposure triggers selection for Cebpa-knockout multipotent hematopoietic progenitors. J Exp Med, 218(6). https://doi.org/10.1084/jem.20200560

Ishida, S., Huang, E., Zuzan, H., Spang, R., Leone, G., West, M., & Nevins, J. R. (2001). Role for E2F in control of both DNA replication and mitotic functions as revealed from DNA microarray analysis. Mol Cell Biol, 21(14), 4684–4699. https://doi.org/10.1128/MCB.21.14.4684-4699.2001

Isshiki, Y., & Iwama, A. (2018). Emerging role of noncanonical polycomb repressive complexes in normal and malignant hematopoiesis. Exp Hematol, 68, 10–14. https://doi.org/10.1016/j.exphem.2018.10.008

Isshiki, Y., Nakajima-Takagi, Y., Oshima, M., Aoyama, K., Rizk, M., Kurosawa, S., Saraya, A., Kondo, T., Sakaida, E., Nakaseko, C., Yokote, K., Koseki, H., & Iwama, A. (2019). KDM2B in polycomb repressive complex 1.1 functions as a tumor suppressor in the initiation of T-cell leukemogenesis. Blood Adv, 3(17), 2537–2549. https://doi.org/10.1182/bloodadvances.2018028522

Iwama, A. (2017). Polycomb repressive complexes in hematological malignancies. Blood, 130(1), 23–29. https://doi.org/10.1182/blood-2017-02-739490

Iwama, A., Oguro, H., Negishi, M., Kato, Y., Morita, Y., Tsukui, H., Ema, H., Kamijo, T., Katoh-Fukui, Y., Koseki, H., van Lohuizen, M., & Nakauchi, H. (2004). Enhanced self-renewal of hematopoietic stem cells mediated by the polycomb gene product Bmi-1. Immunity, 21(6), 843–851. https://doi.org/10.1016/j.immuni.2004.11.004

Kawamoto, T. (2003). Use of a new adhesive film for the preparation of multi-purpose fresh-frozen sections from hard tissues, whole-animals, insects and plants. Arch Histol Cytol, 66(2), 123–143. https://doi.org/10.1679/aohc.66.123

Ku, M., Koche, R. P., Rheinbay, E., Mendenhall, E. M., Endoh, M., Mikkelsen, T. S., Presser, A., Nusbaum, C., Xie, X., Chi, A. S., Adli, M., Kasif, S., Ptaszek, L. M., Cowan, C. A., Lander, E. S., Koseki, H., & Bernstein, B. E. (2008). Genomewide analysis of PRC1 and PRC2 occupancy identifies two classes of bivalent domains. PLoS Genet, 4(10), e1000242. https://doi.org/10.1371/journal.pgen.1000242

Kubosaki, A., Lindgren, G., Tagami, M., Simon, C., Tomaru, Y., Miura, H., Suzuki, T., Arner, E., Forrest, A. R., Irvine, K. M., Schroder, K., Hasegawa, Y., Kanamori-Katayama, M., Rehli, M., Hume, D. A., Kawai, J., Suzuki, M., Suzuki, H., & Hayashizaki, Y. (2010). The combination of gene perturbation assay and ChIP-chip reveals functional direct target genes for IRF8 in THP-1 cells. Mol Immunol, 47(14), 2295–2302. https://doi.org/10.1016/j.molimm.2010.05.289

Liu, Y., Liu, F., Yu, H., Zhao, X., Sashida, G., Deblasio, A., Harr, M., She, Q. B., Chen, Z., Lin, H. K., Di Giandomenico, S., Elf, S. E., Yang, Y., Miyata, Y., Huang, G., Menendez, S., Mellinghoff, I. K., Rosen, N., Pandolfi, P. P., … Nimer, S. D. (2012). Akt phosphorylates the transcriptional repressor bmi1 to block its effects on the tumor-suppressing ink4a-arf locus. Sci Signal, 5(247), ra77. https://doi.org/10.1126/scisignal.2003199

Loke, J., Chin, P. S., Keane, P., Pickin, A., Assi, S. A., Ptasinska, A., Imperato, M. R., Cockerill, P. N., & Bonifer, C. (2018). C/EBPα overrides epigenetic reprogramming by oncogenic transcription factors in acute myeloid leukemia. Blood Adv, 2(3), 271–284. https://doi.org/10.1182/bloodadvances.2017012781

Manz, M. G., & Boettcher, S. (2014). Emergency granulopoiesis. Nat Rev Immunol, 14(5), 302–314. https://doi.org/10.1038/nri3660

Masuda, K., Kubagawa, H., Ikawa, T., Chen, C. C., Kakugawa, K., Hattori, M., Kageyama, R., Cooper, M. D., Minato, N., Katsura, Y., & Kawamoto, H. (2005). Prethymic T-cell development defined by the expression of paired immunoglobulin-like receptors. EMBO J, 24(23), 4052–4060. https://doi.org/10.1038/sj.emboj.7600878

Murakami, K., Sasaki, H., Nishiyama, A., Kurotaki, D., Kawase, W., Ban, T., Nakabayashi, J., Kanzaki, S., Sekita, Y., Nakajima, H., Ozato, K., Kimura, T., & Tamura, T. (2021). A RUNX-CBFβ-driven enhancer directs the Irf8 dose-dependent lineage choice between DCs and monocytes. Nat Immunol, 22(3), 301–311. https://doi.org/10.1038/s41590-021-00871-y

Nacerddine, K., Beaudry, J. B., Ginjala, V., Westerman, B., Mattiroli, F., Song, J. Y., van der Poel, H., Ponz, O. B., Pritchard, C., Cornelissen-Steijger, P., Zevenhoven, J., Tanger, E., Sixma, T. K., Ganesan, S., & van Lohuizen, M. (2012). Akt-mediated phosphorylation of Bmi1 modulates its oncogenic potential, E3 ligase activity, and DNA damage repair activity in mouse prostate cancer. J Clin Invest, 122(5), 1920–1932. https://doi.org/10.1172/JCI57477

Nestorowa, S., Hamey, F. K., Pijuan Sala, B., Diamanti, E., Shepherd, M., Laurenti, E., Wilson, N. K., Kent, D. G., & Göttgens, B. (2016). A single-cell resolution map of mouse hematopoietic stem and progenitor cell differentiation. Blood, 128(8), e20–31. https://doi.org/10.1182/blood-2016-05-716480

Oguro, H., Iwama, A., Morita, Y., Kamijo, T., van Lohuizen, M., & Nakauchi, H. (2006). Differential impact of Ink4a and Arf on hematopoietic stem cells and their bone marrow microenvironment in Bmi1-deficient mice. J Exp Med, 203(10), 2247–2253. https://doi.org/10.1084/jem.20052477

Oguro, H., Yuan, J., Ichikawa, H., Ikawa, T., Yamazaki, S., Kawamoto, H., Nakauchi, H., & Iwama, A. (2010). Poised lineage specification in multipotential hematopoietic stem and progenitor cells by the polycomb protein Bmi1. Cell Stem Cell, 6(3), 279–286. https://doi.org/10.1016/j.stem.2010.01.005

Park, I. K., Qian, D., Kiel, M., Becker, M. W., Pihalja, M., Weissman, I. L., Morrison, S. J., & Clarke, M. F. (2003). Bmi-1 is required for maintenance of adult self-renewing haematopoietic stem cells. Nature, 423(6937), 302–305. https://doi.org/10.1038/nature01587

Pietras, E. M., Mirantes-Barbeito, C., Fong, S., Loeffler, D., Kovtonyuk, L. V., Zhang, S., Lakshminarasimhan, R., Chin, C. P., Techner, J. M., Will, B., Nerlov, C., Steidl, U., Manz, M. G., Schroeder, T., & Passegué, E. (2016). Chronic interleukin-1 exposure drives haematopoietic stem cells towards precocious myeloid differentiation at the expense of self-renewal. Nat Cell Biol, 18(6), 607–618. https://doi.org/10.1038/ncb3346

Pietras, E. M., Reynaud, D., Kang, Y. A., Carlin, D., Calero-Nieto, F. J., Leavitt, A. D., Stuart, J. M., Göttgens, B., & Passegué, E. (2015). Functionally Distinct Subsets of Lineage-Biased Multipotent Progenitors Control Blood Production in Normal and Regenerative Conditions. Cell Stem Cell, 17(1), 35–46. https://doi.org/10.1016/j.stem.2015.05.003

Piunti, A., & Shilatifard, A. (2021). The roles of Polycomb repressive complexes in mammalian development and cancer. Nat Rev Mol Cell Biol, 22(5), 326–345. https://doi.org/10.1038/s41580-021-00341-1

Rosenbauer, F., & Tenen, D. G. (2007). Transcription factors in myeloid development: balancing differentiation with transformation. Nat Rev Immunol, 7(2), 105–117. https://doi.org/10.1038/nri2024

Ross, K., Sedello, A. K., Todd, G. P., Paszkowski-Rogacz, M., Bird, A. W., Ding, L., Grinenko, T., Behrens, K., Hubner, N., Mann, M., Waskow, C., Stocking, C., & Buchholz, F. (2012). Polycomb group ring finger 1 cooperates with Runx1 in regulating differentiation and self-renewal of hematopoietic cells. Blood, 119(18), 4152–4161. https://doi.org/10.1182/blood-2011-09-382390

Sashida, G., & Iwama, A. (2012). Epigenetic regulation of hematopoiesis. Int J Hematol, 96(4), 405–412. https://doi.org/10.1007/s12185-012-1183-x

Sawai, C. M., Babovic, S., Upadhaya, S., Knapp, D. J. H. F., Lavin, Y., Lau, C. M., Goloborodko, A., Feng, J., Fujisaki, J., Ding, L., Mirny, L. A., Merad, M., Eaves, C. J., & Reizis, B. (2016). Hematopoietic Stem Cells Are the Major Source of Multilineage Hematopoiesis in Adult Animals. Immunity, 45(3), 597–609. https://doi.org/10.1016/j.immuni.2016.08.007

Säwen, P., Eldeeb, M., Erlandsson, E., Kristiansen, T. A., Laterza, C., Kokaia, Z., Karlsson, G., Yuan, J., Soneji, S., Mandal, P. K., Rossi, D. J., & Bryder, D. (2018). Murine HSCs contribute actively to native hematopoiesis but with reduced differentiation capacity upon aging. Elife, 7. https://doi.org/10.7554/eLife.41258

Shinoda, D., Nakajima-Takagi, Y., Oshima, M., Koide, S., Aoyama, K., Saraya, A., Harada, H., Rahmutulla, B., Kaneda, A., Yamaguchi, K., Furukawa, Y., Koseki, H., Shimoda, K., Tanaka, T., Sashida, G., & Iwama, A. (2022). Insufficiency of non- canonical PRC1 synergizes with JAK2V617F in the development of myelofibrosis. Leukemia, 36(2), 452–463. https://doi.org/10.1038/s41375-021-01402-2

Shooshtarizadeh, P., Helness, A., Vadnais, C., Brouwer, N., Beauchemin, H., Chen, R., Bagci, H., Staal, F. J. T., Coté, J. F., & Möröy, T. (2019). Gfi1b regulates the level of Wnt/β-catenin signaling in hematopoietic stem cells and megakaryocytes. Nat Commun, 10(1), 1270. https://doi.org/10.1038/s41467-019-09273-z

Si, S., Nakajima-Takagi, Y., Aoyama, K., Oshima, M., Saraya, A., Sugishita, H., Nakayama, M., Ishikura, T., Koseki, H., & Iwama, A. (2016). Loss of Pcgf5 Affects Global H2A Monoubiquitination but Not the Function of Hematopoietic Stem and Progenitor Cells. PLoS One, 11(5), e0154561. https://doi.org/10.1371/journal.pone.0154561

Slomiany, B. A., D’Arigo, K. L., Kelly, M. M., & Kurtz, D. T. (2000). C/EBPalpha inhibits cell growth via direct repression of E2F-DP-mediated transcription. Mol Cell Biol, 20(16), 5986–5997. https://doi.org/10.1128/MCB.20.16.5986-5997.2000

Sonntag, R., Giebeler, N., Nevzorova, Y. A., Bangen, J. M., Fahrenkamp, D., Lambertz, D., Haas, U., Hu, W., Gassler, N., Cubero, F. J., Müller-Newen, G., Abdallah, A. T., Weiskirchen, R., Ticconi, F., Costa, I. G., Barbacid, M., Trautwein, C., & Liedtke, C. (2018). Cyclin E1 and cyclin-dependent kinase 2 are critical for initiation, but not for progression of hepatocellular carcinoma. Proc Natl Acad Sci U S A, 115(37), 9282–9287. https://doi.org/10.1073/pnas.1807155115

Sun, J., Ramos, A., Chapman, B., Johnnidis, J. B., Le, L., Ho, Y. J., Klein, A., Hofmann, O., & Camargo, F. D. (2014). Clonal dynamics of native haematopoiesis. Nature, 514(7522), 322–327. https://doi.org/10.1038/nature13824

Takayama, N., Murison, A., Takayanagi, S. I., Arlidge, C., Zhou, S., Garcia-Prat, L., Chan-Seng-Yue, M., Zandi, S., Gan, O. I., Boutzen, H., Kaufmann, K. B., Trotman-Grant, A., Schoof, E., Kron, K., Díaz, N., Lee, J. J. Y., Medina, T., De Carvalho, D. D., Taylor, M. D., … Lupien, M. (2021). The Transition from Quiescent to Activated States in Human Hematopoietic Stem Cells Is Governed by Dynamic 3D Genome Reorganization. Cell Stem Cell, 28(3), 488–501.e410. https://doi.org/10.1016/j.stem.2020.11.001

Tara, S., Isshiki, Y., Nakajima-Takagi, Y., Oshima, M., Aoyama, K., Tanaka, T., Shinoda, D., Koide, S., Saraya, A., Miyagi, S., Manabe, I., Matsui, H., Koseki, H., Bardwell, V. J., & Iwama, A. (2018). Bcor insufficiency promotes initiation and progression of myelodysplastic syndrome. Blood, 132(23), 2470–2483. https://doi.org/10.1182/blood-2018-01-827964

Trumpp, A., Essers, M., & Wilson, A. (2010). Awakening dormant haematopoietic stem cells. Nat Rev Immunol, 10(3), 201–209. https://doi.org/10.1038/nri2726

Voncken, J. W., Niessen, H., Neufeld, B., Rennefahrt, U., Dahlmans, V., Kubben, N., Holzer, B., Ludwig, S., & Rapp, U. R. (2005). MAPKAP kinase 3pK phosphorylates and regulates chromatin association of the polycomb group protein Bmi1. J Biol Chem, 280(7), 5178–5187. https://doi.org/10.1074/jbc.M407155200

Wang, L., Brown, J. L., Cao, R., Zhang, Y., Kassis, J. A., & Jones, R. S. (2004). Hierarchical recruitment of polycomb group silencing complexes. Mol Cell, 14(5), 637–646. https://doi.org/10.1016/j.molcel.2004.05.009

Wang, Y., Krivtsov, A. V., Sinha, A. U., North, T. E., Goessling, W., Feng, Z., Zon, L. I., & Armstrong, S. A. (2010). The Wnt/beta-catenin pathway is required for the development of leukemia stem cells in AML. Science, 327(5973), 1650–1653. https://doi.org/10.1126/science.1186624

Xie, H., Ye, M., Feng, R., & Graf, T. (2004). Stepwise reprogramming of B cells into macrophages. Cell, 117(5), 663–676. https://doi.org/10.1016/s0092-8674(04)00419-2

Ye, M., Zhang, H., Amabile, G., Yang, H., Staber, P. B., Zhang, P., Levantini, E., Alberich- Jordà, M., Zhang, J., Kawasaki, A., & Tenen, D. G. (2013). C/EBPa controls acquisition and maintenance of adult haematopoietic stem cell quiescence. Nat Cell Biol, 15(4), 385–394. https://doi.org/10.1038/ncb2698

Zhang, D. E., Zhang, P., Wang, N. D., Hetherington, C. J., Darlington, G. J., & Tenen, D. G. (1997). Absence of granulocyte colony-stimulating factor signaling and neutrophil development in CCAAT enhancer binding protein alpha-deficient mice. Proc Natl Acad Sci U S A, 94(2), 569–574. https://doi.org/10.1073/pnas.94.2.569

Zhang, P., Iwasaki-Arai, J., Iwasaki, H., Fenyus, M. L., Dayaram, T., Owens, B. M., Shigematsu, H., Levantini, E., Huettner, C. S., Lekstrom-Himes, J. A., Akashi, K., & Tenen, D. G. (2004). Enhancement of hematopoietic stem cell repopulating capacity and self-renewal in the absence of the transcription factor C/EBP alpha. Immunity, 21(6), 853–863. https://doi.org/10.1016/j.immuni.2004.11.006

Zhao, J. L., & Baltimore, D. (2015). Regulation of stress-induced hematopoiesis. Curr Opin Hematol, 22(4), 286–292. https://doi.org/10.1097/MOH.0000000000000149

